# Period five indicates autopoiesis

**DOI:** 10.64898/2026.07.29.741398

**Authors:** Peng Bao, Min Qiu

**Affiliations:** Institute of Urban Environment, Chinese Academy of Sciences, Xiamen 361021, China; University of Chinese Academy of Sciences, Beijing 100049, China; College of Life Sciences, Fujian Agriculture and Forestry University, Fuzhou 350002, China

**Author notes:** **Corresponding Author:** Institute of Urban Environment, Chinese Academy of Sciences, Xiamen 361021, China.

**Keywords:** autopoiesis, weakly reversible realization, membrane compartment, peptide nucleic acids (PNA), linear constraint, period five

## Abstract

A key scientific challenge is to develop a universal theory of life that integrates our biological knowledge with fundamental logical principles. We propose that a "five nodes" principle may govern the origin of life and consistently exist hierarchically within living systems. In our investigation, we explored autocatalytic chemical reaction networks (CRNs) as potential origins for methanotrophy and anoxygenic phototrophy, aiming to validate the "five nodes" principle in the emergence of autopoietic systems. Our research revealed the emergence of autocatalytic peptides and weakly reversible realizations within the MSA reaction network (composed of CH_4_, SO_4_^2-^/SO_3_^2-^, and NH_4_^+^) as well as in the light-Sammox (sulfurous reduction coupled to anaerobic ammonium oxidation)-driven CRN (composed of HCO_3_^-^, SO_3_^2-^, and NH_4_^+^) under hydrothermal conditions. Furthermore, we identified the possible emergence of three main interdependent components essential for life within the two reaction networks: membrane compartments, peptide nucleic acids (PNA) backbones, and catalytic capacities for energy release reactions. Our findings suggest that non-equilibrium synergy of five bioessential elements (NESFBE) can facilitate proto-energy and material metabolism, thereby enabling diverse scenarios for life’s origin. Importantly, our discovery indicates linear constraints present in these CRNs that contributed to life’s inception, which is the mathematical foundation of the "five nodes" principle. Linear constraints determine both the emergence and self-disintegration of autopoietic systems. We infer a period five existence based on hierarchical structures found within autopoietic systems, and “period five indicates autopoiesis” could be one of universal theory of life.

## 1. Introduction

### 1.1 Theoretical background

A significant long-term scientific challenge that requires attention is the development of a universal theory of life, which integrates our empirical understanding of biology with overarching logical principles [1–3]. To date, efforts to establish principles that are independent of evolved constraints and biochemical materials have not produced comprehensive theories for identifying, quantifying, or creating life. We contend that a universal theory of life should govern the origin of life and consistently exist hierarchically within living systems. Furthermore, unraveling the mystery surrounding the origin of life necessitates the identification of a mathematical physical principle since mathematics offers an all-encompassing framework for elucidating all phenomena.

The early origin of life can be seen as the evolution from the self-organized maximum entropy production and energy seeking chemical reaction network (CRN) (composed of carbon (C), hydrogen (H), oxygen (O), nitrogen (N), and sulfur (S)) to the biochemical network [4]. According to our previous research that HCO_3_^-^, SO_3_^2-^/SO_4_^2-^, NH_4_^+^ and H_2_O were converted into autocatalytic peptides from CRN initiated by sulfurous reduction coupled to anaerobic ammonium oxidation (Sammox) under mild hydrothermal conditions, which was a feasible scenario for peptides evolution and life origin [4], [5]. We therefore propose a “non-equilibrium synergy of five bioessential elements” (NESFBE) hypothesis that a non-equilibrium CRN should at least contain C, H, O, N, and S can lead to the origin of life. Hence, the simultaneous emergence of primordial energy and material metabolism could be feasibly achieved. By using graph theory, we can understand why C, H, O, N, and S are suitable to drive the origin of life. The periodic table of chemical elements can be formalized as a set that includes a system of similarity classes, with each class represented as a hyperedge. Each element within this set holds an order relation, and this structure can be represented as an ordered hypergraph. The hyperedges of bonds of C, H, O, N, and S are the most polarized, giving them dominance over other elements in terms of the number of hyperedges they occupy (Scheme 1) [6]. The C, H, O, N, and S in the NESFBE-CRN plus the initial metallic bio-elements necessary for life, Fe, Mg, Ni, Mo, and W, etc., may form the hypergraph collaborative network. We are keen to understand why the synergy of five bioessential elements can activate primordial energy and facilitate material metabolism. We propose to refer to this phenomenon as the "five nodes" in order to highlight the distinctive role that the number five plays within network systems, suggesting it may represent a mathematical physical principle underlying life origin.

**Scheme 1.**
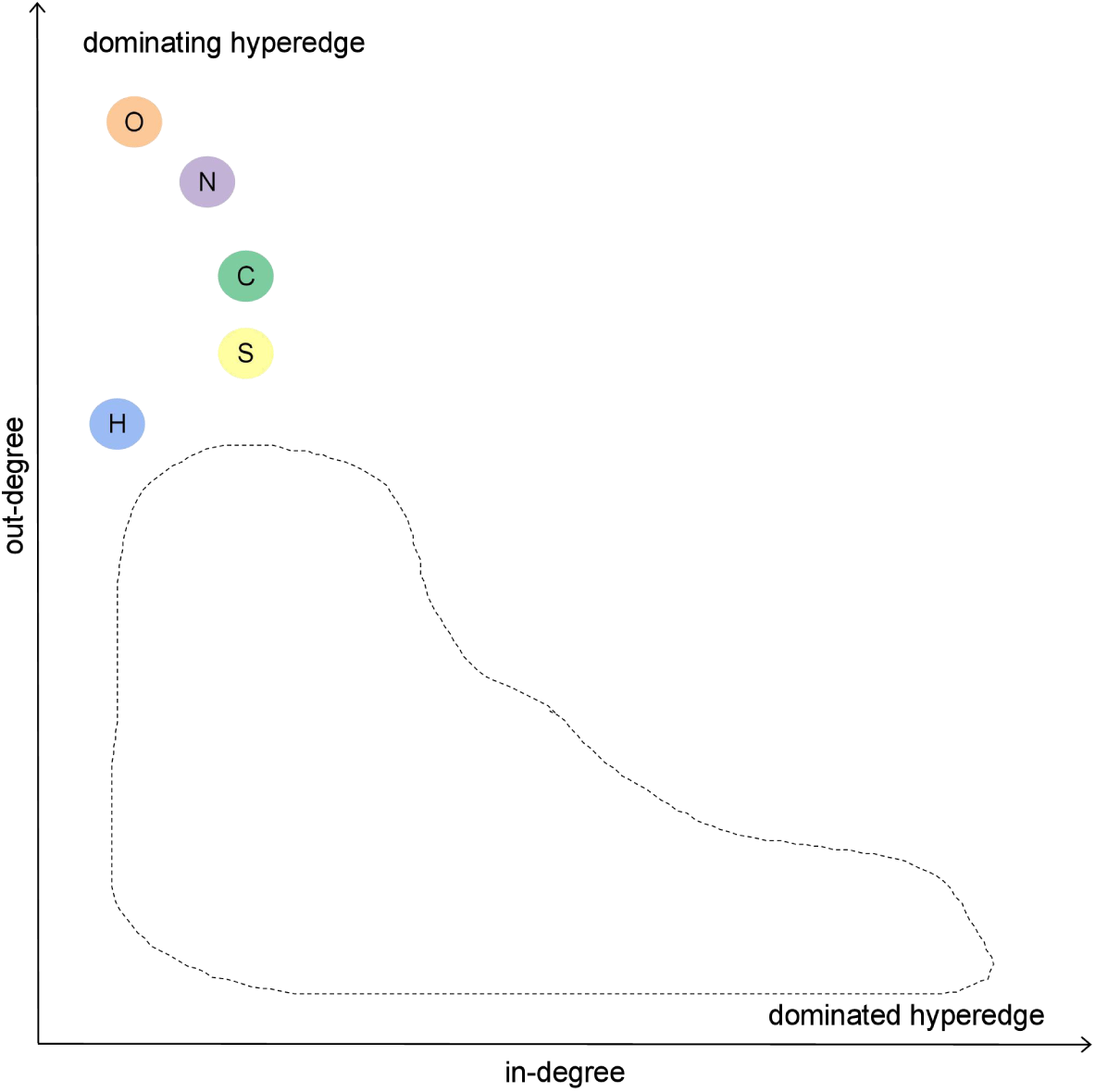
The conceptual model of dominance profile for C, H, O, N, and S hyperedges of single covalent bonds (modified from Ref [6]). Other elements are within the dashed box and not displayed.

Further, we noticed that the "five nodes" principle is evident in several complex network studies. One such study is Sampson’s monk network, which examines the social relationships among monks in a secluded American monastery, as documented by Sampson in 1969 [7]. In this study, it was found that the minimum controlling set consists of five nodes, indicating that only five monks are required to control the centrality of the social network [8]. Youssef and Scoglio (2013) proposed a solution to the biological epidemic mitigation problem in a networked population, where they found that only a network with five nodes could yield optimal results [9]. For dynamic resource optimization without seed selection, Kandhway and Kuri (2017) discovered that five groups are sufficient to capture the disparity in centrality measures for implementing dynamic controls [10]. However, joint allocation consistently achieves better results as the number of groups increases. Nevertheless, the percentage improvement diminishes rapidly with the addition of more groups. Albert et al (2000) reported that both exponential and scale-free networks, consisting of 130 nodes and 215 links, exhibit five highly connected nodes [11]. However, in the exponential network, only 27% of the nodes are reached by these five most connected nodes, whereas in the scale-free network, more than 60% of the nodes are reached [11]. We have discovered that the "five nodes" principle may be intricately linked to the emergence and self-maintenance of complex systems. Consequently, our intrinsic curiosity is directed towards understanding the fundamental principles underlying the emergence and evolution of autopoietic systems, which are characterized by self-production and self-maintenance [12]. Among these topics, the origin of life emerges as a particularly significant area of inquiry.

The origins of primordial metabolism, particularly methanogenesis and anoxygenic phototrophy, represent fundamental biological questions that have long awaited resolution. We align with the perspective that life has emerged multiple times on Earth and that various forms of extant life coexist across a diverse array of physical substrates [3]. In this study, we aim to investigate the plausibility of autocatalytic primordial chemical reaction networks (CRNs) as potential origin scenarios for methanogenesis (CH_4_, SO_3_^2-^/SO_3_^2-^, and NH_4_^+^) and anoxygenic phototrophy (HCO_3_^-^, SO_3_^2-^, NH_4_^+^, and Light radiation), within the framework of NESFBE. This approach seeks to verify the feasibility of the "five nodes" hypothesis. We will present comprehensive background information regarding the hypothetical origins of methanogenesis and anoxygenic phototrophy in the following sections.

### 1.2 Methanogenesis origin and theoretical hypothesis

Methane has been proposed as an alternative to carbon dioxide for ancient carbon fixation pathways dating back to the origin of life [13], [14]. Methanogenesis, sulfur reduction, sulfate reduction, and anoxygenic photosynthesis are believed to have appeared between 3.8 and 3.4 billion years ago [15]. The denitrifying methanotrophic acetogenic pathway, which is a type of reverse methanogenesis, may have emerged earlier than traditional methanogenesis under hydrothermal conditions (Eqs 1 and 2) [14], [16].

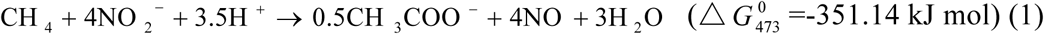

[13], [17]

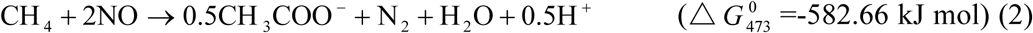

[13], [17]

It is plausible to hypothesize that methane can serve not only as a fuel but also as a readily available source of reduced carbon to drive CRN towards proto-metabolic network, provided that there is a continuous supply of nitrite/nitrate and sulfurous species in the CRN. We infer that a CRN that includes methane, sulfurous species, and ammonium (termed MSA reaction network) can fulfill the sulfur requirements for NESFBE-CRN hypothesis and provide nitrite for primordial denitrifying methanotrophy via the Sammox reaction (Eq 3) [4], [18].

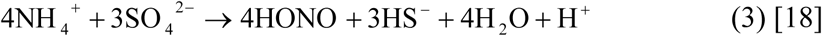

### 1.3 Anoxygenic phototrophy origin and theoretical hypothesis

The origin of anoxygenic photosynthesis has been a fundamental biological question for decades. Phylogenetic analysis of Mg-tetrapyrrole pigments, including chlorophyll and bacteriochlorophyll biosynthesis genes, suggests that most anoxygenic phototrophs are ancestral [15], [19]. Additionally, purple bacteria may contain the most ancestral form of this pigment biosynthesis pathway [15], [19]. Purple bacteria are capable of performing anaerobic photosynthesis through the oxidation of reduced sulfur compounds, which is facilitated by sulfite oxidoreductase and/or thiosulfate-oxidizing (Sox) multi-enzyme complex [20]. Additionally, some purple sulfur bacteria can utilize externally available sulfite as a photosynthetic electron donor through the action of sulfite-oxidizing enzyme (SoeABC), a membrane-bound polysulfide reductase-like iron-sulfur molybdoprotein [21]. Recent research has demonstrated that photoredox cycling, induced by UV irradiation of ferrocyanide with stoichiometric sulfite, can effectively drive the reductive homologation of hydrogen cyanide (HCN) to produce simple sugars and precursors of hydroxy acids and amino acids [22]. Sammox-driven CRNs mentioned in NESFBE hypothesis can meet those elements: continuous supply of hydrogen cyanide and redox cycling of sulfite [4], provided light stimulates. Fortunately, light is abundant in the high-temperature hydrothermal vents where these reactions are believed to occur [23], [24]. At both black smokers and flange pools, light emission is highest at long wavelengths (>700 nm) due to thermal radiation. In addition, black smokers emit time-varying radiation in the visible region (400-650 nm) caused by mechanisms related to turbulence, mixing, or precipitation [23–26]. The isolation and cultivation of obligately photoautotrophic bacteria from hydrothermal vents samples have raised the possibility that low-light emission at hydrothermal vents could support anoxygenic phototrophy [27], [28].

The possibility of a light-driven origin of anoxygenic photosynthesis in a CRN at hydrothermal vents is a promising scenario for the origin of life, compatible with NESFBE hypothesis. We noticed an overlap between light-driven CRN (including HCO_3_^-^, SO_3_^2-^, NH_4_^+^ and H_2_ O) and Sammox-driven CRN (including HCO_3_^-^, SO_3_^2-^/SO_4_^2-^, NH_4_^+^ and H_2_O), an earlier reported life origin scenario [4]. This overlap suggests that Sammox-driven CRN can provide a steady supply of hydrogen cyanide for light-driven CRN, further supporting the feasibility of light-Sammox-driven CRN as anoxygenic phototrophy origin scenario. In summery, Sammox-driven CRN overlap with both MSA reaction network and light-Sammox-driven CRN (Scheme 2).

**Scheme 2.**
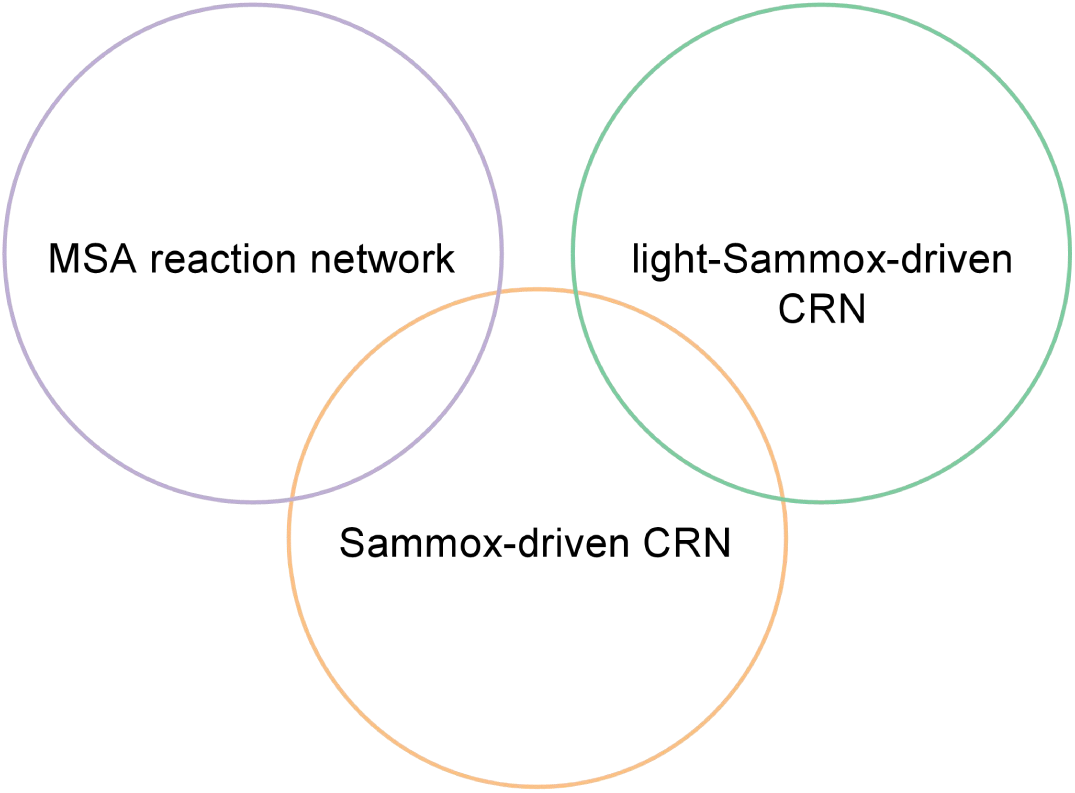
The overlapping relationship of Sammox-driven CRN, MSA reaction network, and light-Sammox-driven CRN.

### 1.4 verification roadmap for the "five nodes" principle

Autocatalysis, defined as the capacity of chemical systems to generate more of themselves, is a fundamental characteristic of living systems [29]. It plays a pivotal role in various biological processes, including metabolism, reproduction, and evolution. The study conducted by Blokhuis et al. (2020) revealed that autocatalytic networks exhibit five distinct motifs serving as autocatalytic cores [30]. A critical aspect that should not be overlooked is the necessity for weak reversibility as an essential feature for establishing a sustainable autocatalytic CRN [31]. We have indeed observed this phenomenon in our previous research in Sammox-driven CRN [4]. In subsequent investigations, we further elaborated on the presence of weak reversibility within Sammox-driven CRNs utilizing nitrogen isotope-labeled glycine [5].

In exploring the origin of life, it is imperative to address how the three main interdependent components metabolic machinery, membrane compartments, and template/genetic mechanisms emerged [32]. Consequently, we aim to employ two non-equilibrium chemical reaction systems (MSA reaction network and light-Sammox-driven CRN) composed of five bioessential elements to demonstrate both the spontaneous emergence of life-like autocatalytic behavior and weak reversibility. Building upon this foundation, we will also investigate the potential emergence of metabolic machinery, membrane compartments, and template/genetic mechanisms as circumstantial evidence supporting the "five nodes" principle. Our primary objectives are outlined as follows:

1. Life-like autocatalytic behavior: Investigating the formation of autocatalytic peptides/products and characterizing autocatalysis within both the MSA reaction network and light-driven Sammox CRN under hydrothermal conditions.
2. Weakly reversible realizations: Examining reverse reactions in both the MSA reaction network and light-driven Sammox CRN under hydrothermal conditions.
3. Emergence of three main interdependent components: Focusing on membrane compartments and peptide nucleic acids (PNA) backbones, and the catalytic capacity for energy release reactions in these two CRNs.

We will attempt to prove that the "five nodes" principle is a universal theory of life by achieving the aforementioned findings. Ultimately, our objective is to elucidate the mathematical logic associated with the number five within the context of autopoietic systems and to further clarify the underlying mathematical and physical principles that govern the emergence of life.

## 2. Results and discussion

### 2.1 Peptides formation in MAS reaction network and light-Sammox-driven CRN

#### 2.1.1 Peptides formation in MSA reaction network

Our analysis revealed that formate and acetate are the primary organic carbon products generated by the MSA reaction network (Figure 1). Notably, the concentration of acetate was found to be higher than that of formate (Figure 1). Additionally, we observed the consumption of methane, sulfurous, and ammonium in the MSA reaction network (Figure 1). Thiosulfate and sulfate were identified as the main intermediates in the sulfur cycle [4] of the MSA reaction network.

**Figure 1.**
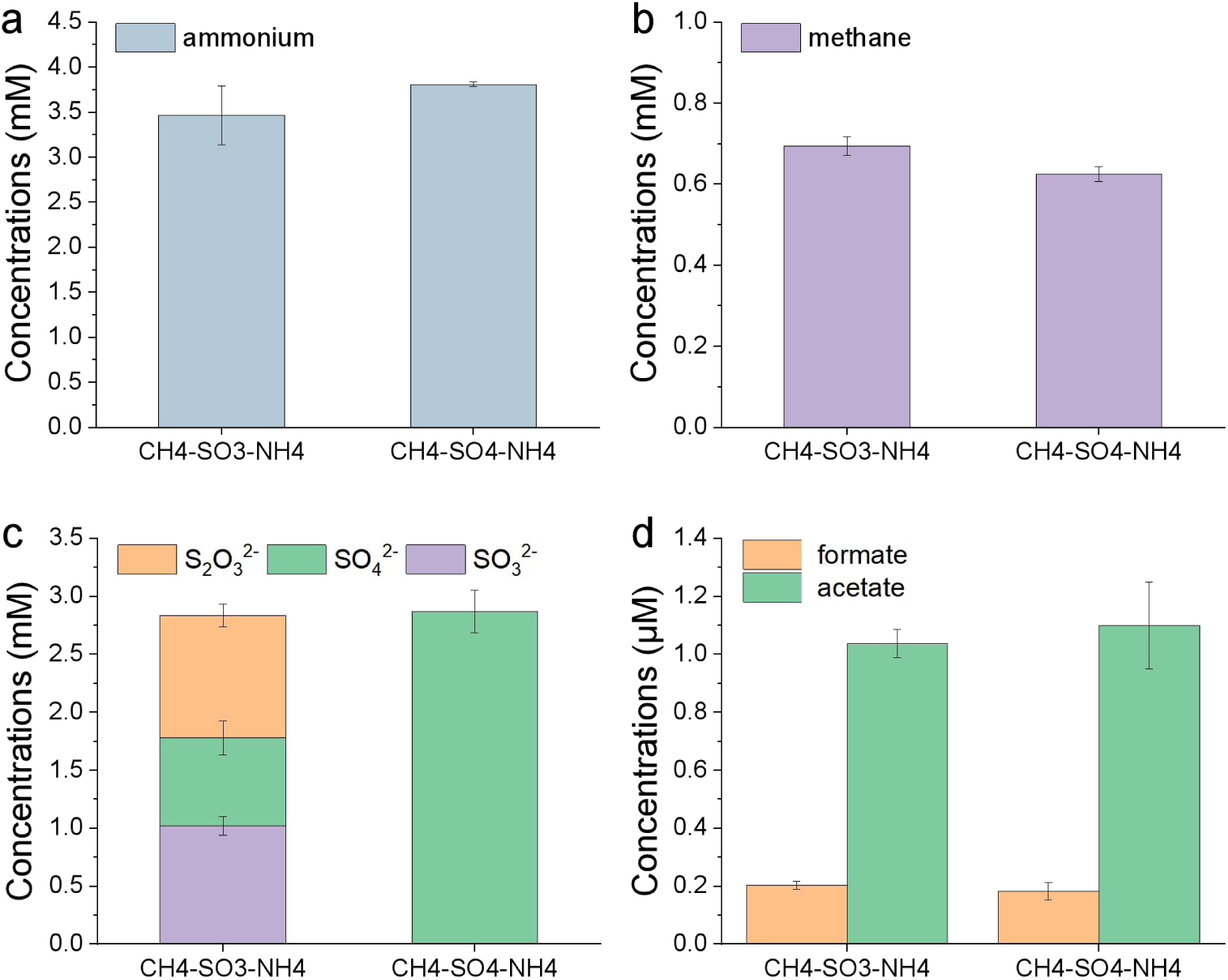
Substrates and primary products transformation in MSA reaction network. Treatments were as follows (CH_4_-SO_3_-NH_4_, sulfite-fueled MSA reaction network; CH_4_-SO_4_-NH_4_, sulfate-fueled MSA reaction network) from left to right. Error bars represent standard deviations of three replicates.

Our study detected peptides as a product of MSA reaction network, as shown in Table 1 and Figure 2 a-f. Table 1 presents selected peptides identified from the MSA reaction network, while Figure 2 a-f shows representative MS/MS spectra of the identified peptides. These peptides were composed of 13 proteinogenic amino acids, namely alanine, glycine, valine, leucine, serine, aspartate, lysine, glutamate, tyrosine, threonine, proline, phenylalanine, and isoleucine (Table 1; Figure 2 a-f). Total ion flow chromatography of peptides and chromatogram of amino acids were shown in SI.

**Figure 2.**
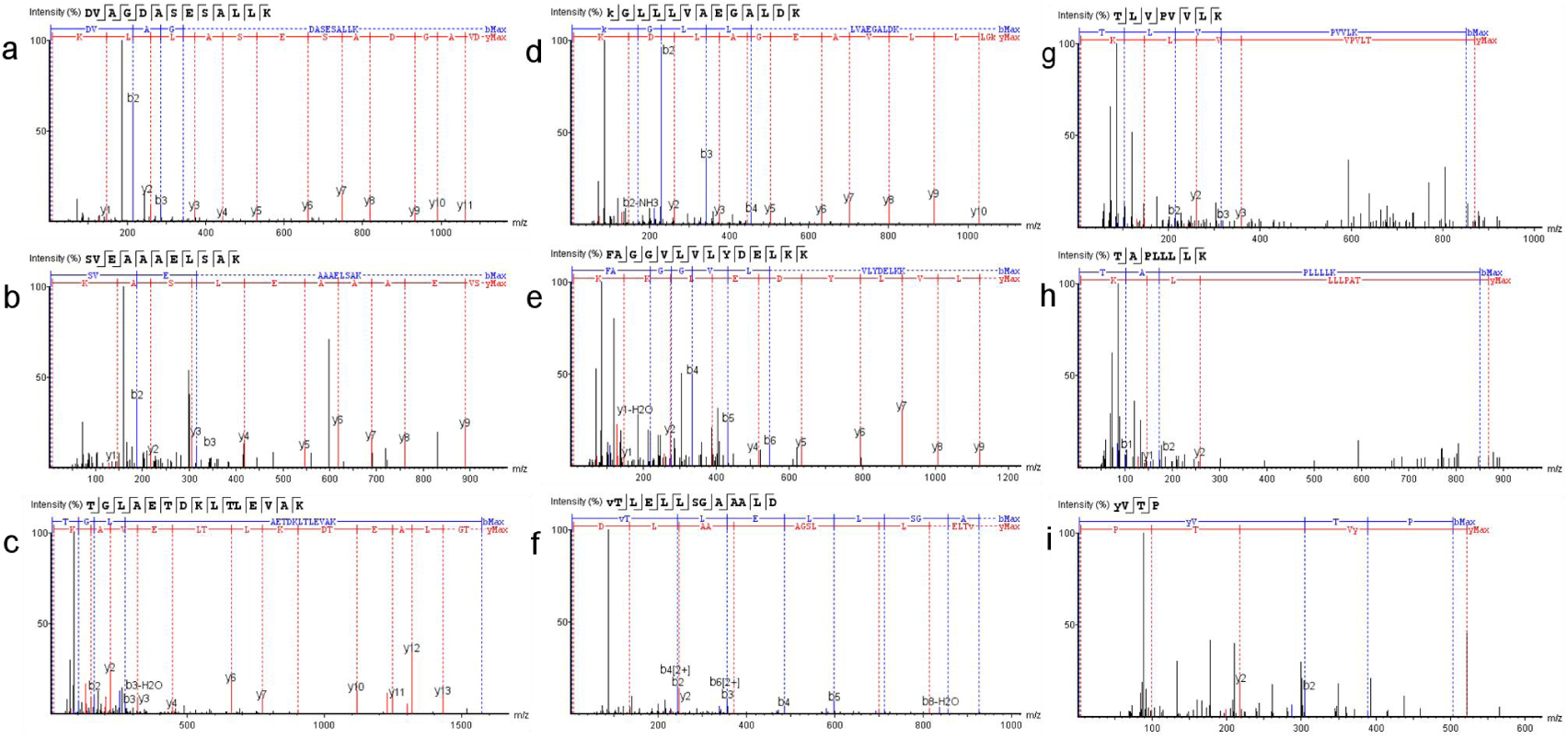
MS/MS spectra of selected peptides in MSA reaction network and light-Sammox-driven CRN corresponding to table 1. All experiences were conducted in positive electrospray ionization mode. The spectra of selected three peptides a–c were from sulfite-fueled MSA reaction network. The spectra of selected three peptides d–f were from sulfate-fueled MSA reaction network; The spectra of selected three peptides g-i were from light-Sammox-driven CRN. The left side of all peptides sequences in the figure is N-terminal and the right side is C-terminal.

**Table 1.** Identified selected peptides in MSA reaction network and light-Sammox-driven CRN. Sulfi-MSA, peptides from sulfite-fueled MSA reaction network; Sulfa-MSA, peptides from sulfate-fueled MSA reaction network; LS-CRN, peptides from light-Sammox-driven CRN.

| Peptides ID | Denovo peptides | m/z | Area | Denovo Score | ppm |
| --- | --- | --- | --- | --- | --- |
| Sulfi-MSA-1 | DVAGDASESALLK | 638.3254 | 3.43E+0.7 | 99 | 0.7 |
| Sulfi-MSA-2 | SVEAAAELSAK | 538.2853 | 1.55E+0.6 | 98 | 0.3 |
| Sulfi-MSA-3 | TGLAETDKLTLEVAK | 530.2988 | 1.86E+0.7 | 97 | 1.8 |
| Sulfa-MSA-1 | K(+42.01)GLLLVAEGAL<br>DK | 684.9088 | 2.39E+0.6 | 96 | 0.5 |
| Sulfa-MSA-2 | FAGGVLVLYDELKK | 776.442 | 1.75E+0.7 | 95 | -1 |
| Sulfa-MSA-3 | V(+42.01)TLELLSGAAA<br>LD | 657.8613 | 1.13E+0.7 | 94 | 0.3 |
| LS-CRN-1 | TLVPVVLK | 868.5806 | 5.27E+0.5 | 94 | -6.9 |
| LS-CRN-2 | TAPLLLLK | 868.588 | 2.37E+0.5 | 91 | 1.7 |
| LS-CRN-3 | Y(+42.01)VTP | 521.2581 | 3.09E+0.5 | 90 | -4.8 |

The oxidation of methane with nitrogen oxide compounds is highly thermodynamically favorable, as demonstrated by equations 1 and 2. This process can result in the formation of carboxylic and keto acids, as noted by Marakushev and Belonogova (2019) [13]. Our analysis detected the presence of acetate, which aligns with the chemical equations 1 and 2. Additionally, we observed the presence of formate, which may correspond to equation 4.

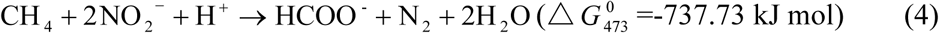

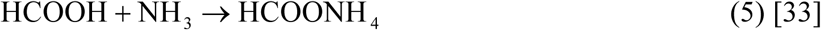

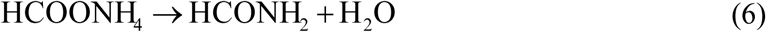

The MSA reaction network might follow the HCOOH→HCOONH_4_→HCONH_4_→HCN route for synthesizing amino acids [4], [34] (Eqs 5 and 6). In addition, HCOONH_4_ may be both hydrogen and nitrogen sources for the reductive amination of α-keto acids to synthesis amino acids [4], [33]. In hydrothermal conditions, a portion of formate breaks down to release carbon dioxide. Additionally, the interaction of hydrogen sulfide and carbon dioxide under hydrothermal conditions can generate carbon disulfide (CS_2_) and carbonyl sulfide (COS), which have been shown to promote peptide bond formation [35–37].

#### 2.1.2 Peptides formation in light-Sammox-driven CRN

Our study revealed that formate is the primary organic carbon product of light-Sammox-driven CRN, as demonstrated in Figure 3. Additionally, we detected peptides as a product of this CRN, as shown in Figure 2 g-i. Table 1 presents a selection of identified peptides produced from light-Sammox-driven CRN, and Figure 2 g-i provides representative MS/MS spectra of these peptides. Table 1 and Figure 2 g-i show that the peptides consist of 13 proteinogenic amino acids, namely alanine, glycine, valine, leucine, serine, aspartate, lysine, glutamate, tyrosine, threonine, proline, phenylalanine, and isoleucine. In the light-Sammox-driven CRN (Figure 3), sulfurous and ammonium are consumed, with thiosulfate and sulfate serving as the main intermediates in the sulfur cycle. Total ion flow chromatography of peptides and chromatogram of amino acids were shown in supporting information.

**Figure 3.**
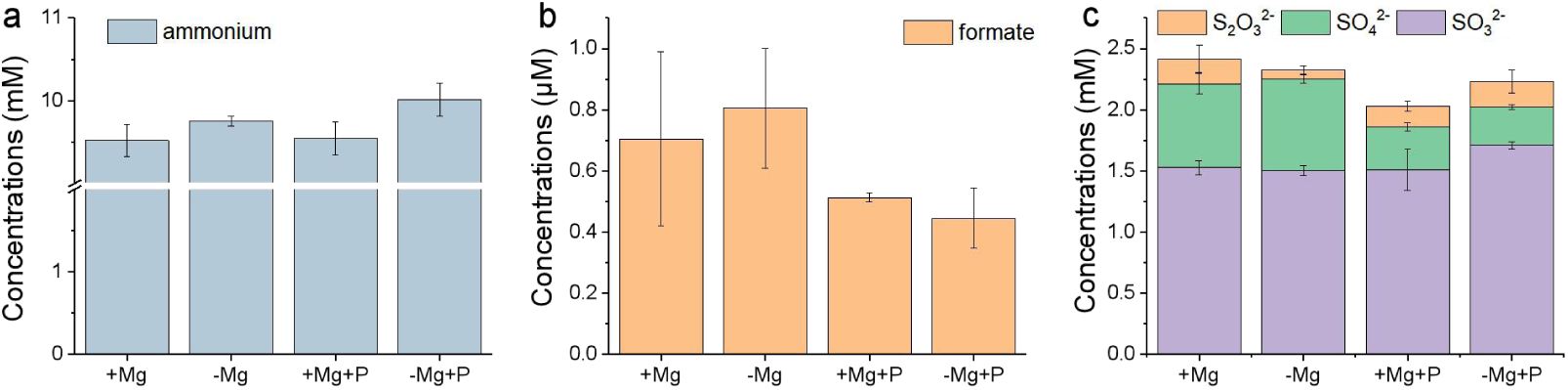
Substrates and primary products transformation in light-Sammox-driven CRN. Treatments were as follows (+Mg, light-Sammox-driven CRN with Mg^2+^; -Mg, light-Sammox-driven CRN; +Mg+P, light-Sammox-driven CRN with Mg^2+^ + products from light-Sammox-driven CRN with Mg^2+^; -Mg+P, light-Sammox-driven CRN + products from light-Sammox-driven CRN) from left to right. Error bars represent standard deviations of three replicates.

The first step in amino acid synthesis in light-Sammox-driven CRN may involve a photochemical reductive homologation of hydrogen cyanide (HCN) by sulfite. Sulfite is activated early in the reaction (as shown in Eq 7) and then reacts/catalyzes with HCN to generate precursors of hydroxy acids and amino acids [22], [38]. These precursors can be converted to amino acids with the assistance of an activated ammonia donor, HCOONH_4_. In this study, both hydrogen cyanide and HCOONH_4_ are provided by Sammox-driven CRN [4] (Eq 5). Furthermore, Sammox-driven CRN can facilitate amino acid polymerization [4]. Sulfite acts as both a reducing agent and an oxidant in light-Sammox-driven CRN. Figure 4 illustrates the differentiation between light-Sammox-driven CRN and Sammox-driven CRN at 30°C, suggesting the existence of distinct amino acids synthesis pathways. Light-Sammox-driven CRN exhibited greater efficiency in proteinogenic amino acids synthesis compared to Sammox-driven CRN. The results suggest that both photochemical and hydrothermal reactions are essential for peptides generation in light-Sammox-driven CRN.

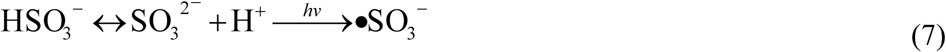

**Figure 4.**
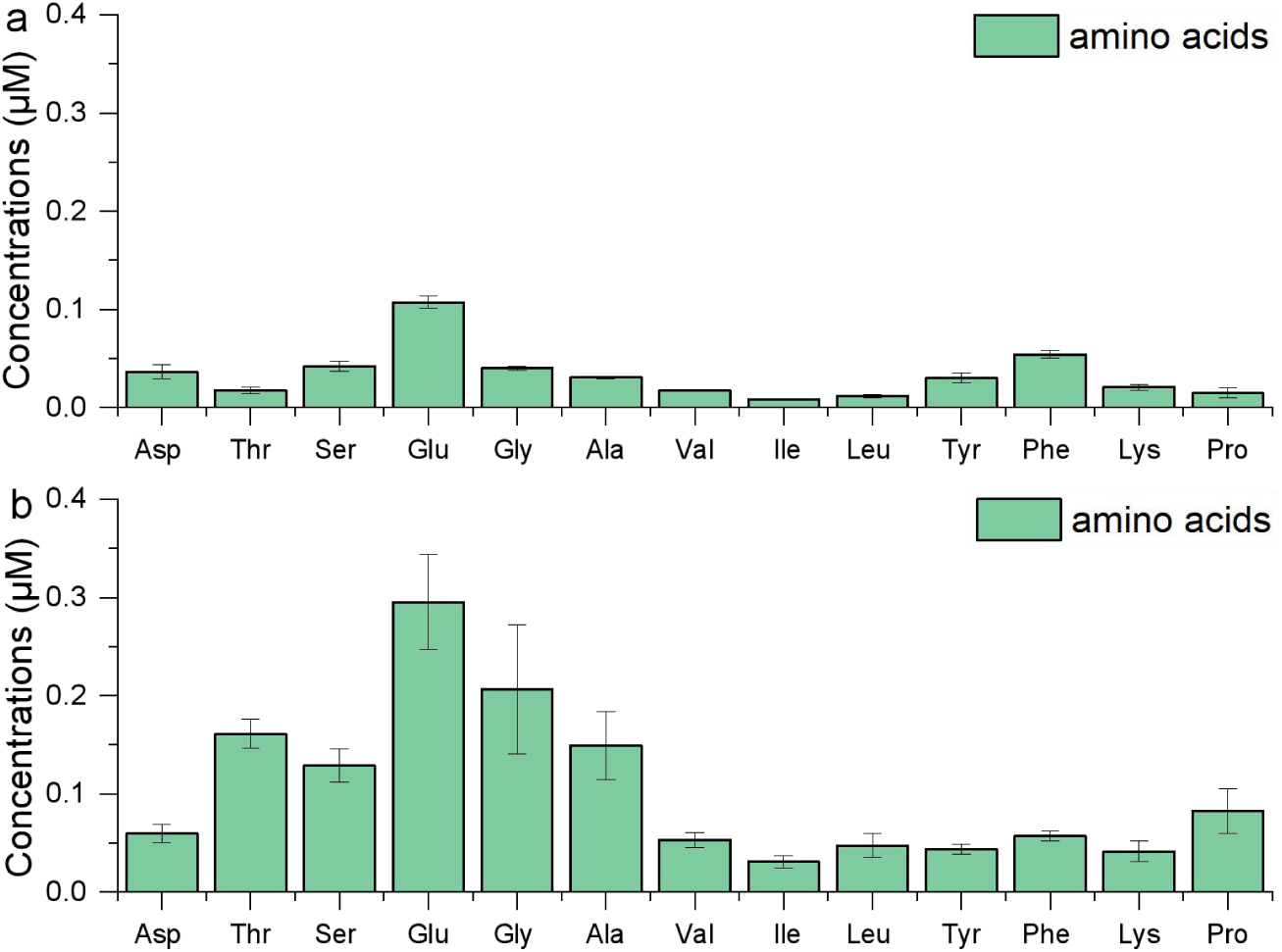
Proteinogenic amino acids generated in Sammox-driven CRN (a) and light-Sammox-driven CRN (b). Error bars represent standard deviations of three replicates.

### 2.2 autocatalysis and weakly reversible realization in MSA reaction network and light-Sammox-driven CRN

#### 2.2.1 autocatalysis in MSA reaction network and light-Sammox-driven CRN

In the experiments outlined in supporting information section D, we conducted a series of investigations to determine if the MSA reaction network exhibits autocatalysis. To achieve this, we utilized MSA reaction network products as a catalyst to catalyze the production of itself. Specifically, we injected a 1.0 mL reaction solution (the first round sulfite/sulfate-fueled MSA reaction network) into a freshly prepared reaction solution (the second round sulfite/sulfate-fueled MSA reaction network) as a potential catalyst. The products generated from both ether sulfite-fueled and sulfate-fueled MSA reaction networks have been found to facilitate the production of formate, acetate, and peptides at 100°C in both sulfite and sulfate-fueled MSA reaction networks (Figure 5).

**Figure 5.**
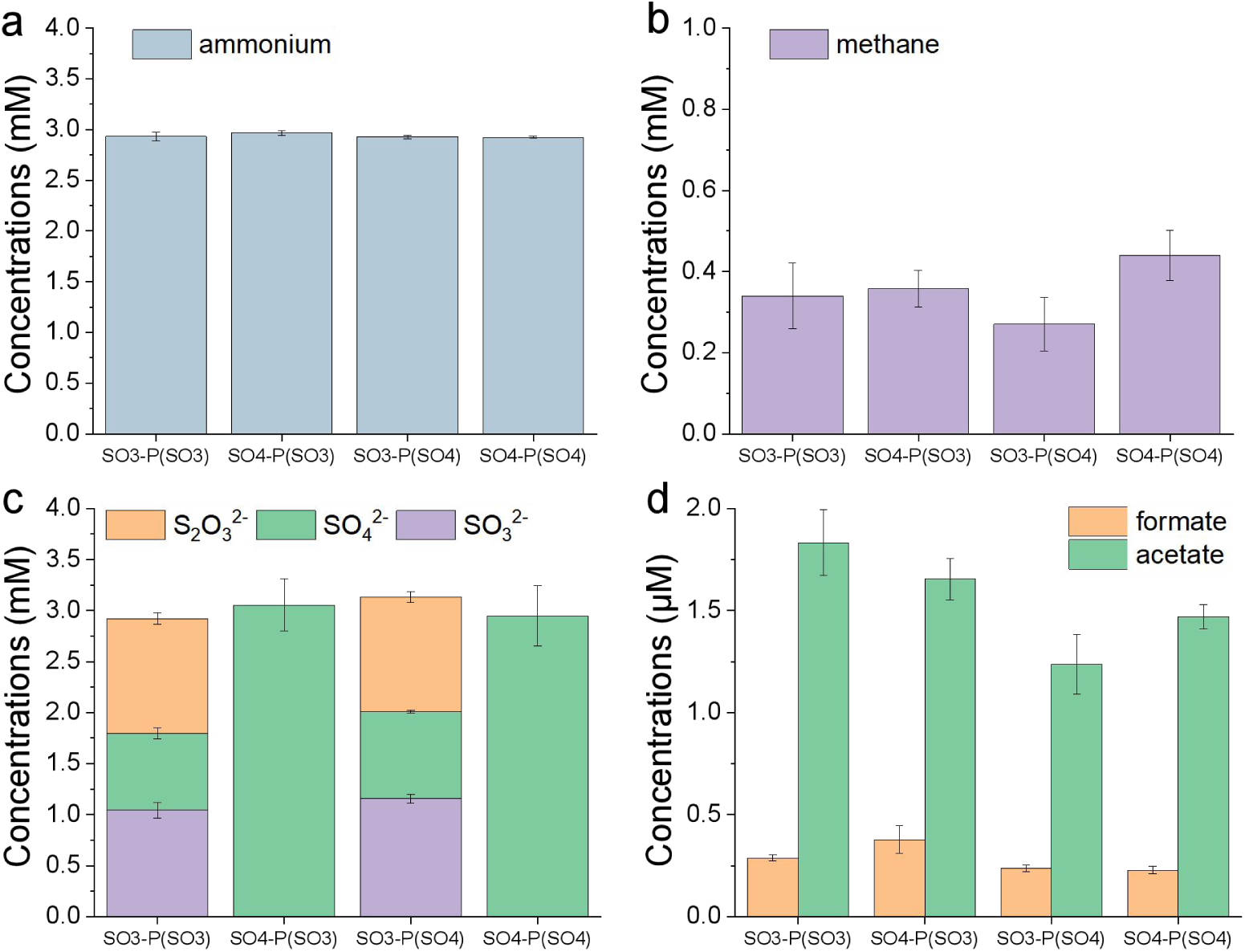
Substrates and primary products transformation in MSA reaction network. From left to right treatments were as follows, SO_3_-P(SO_3_), sulfite-fueled MSA reaction network with addition of catalytic products from sulfite-fueled MSA reaction; SO_4_-P(SO_3_), sulfate-fueled MSA reaction network with addition of catalytic products from sulfite-fueled MSA reaction network; SO_3_-P(SO_4_), sulfite-fueled MSA reaction network with addition of catalytic products from sulfate-fueled MSA reaction network; SO_4_-P(SO_4_), sulfate-fueled MSA reaction network with addition of catalytic products from sulfate-fueled MSA reaction network. Error bars represent standard deviations of three replicates.

The peptides generated in the sulfite-fueled MSA reaction network with catalytic products from the sulfite/sulfate-fueled MSA reaction treatment groups appear to be slightly less than those in the sulfite-fueled MSA reaction network (Figure 6a, b, and c). However, the presence of additional methionine, histidine, and arginine in the sulfite-fueled MSA reaction network with catalytic products from the sulfite/sulfate-fueled MSA reaction treatment groups (Figure 6b and c) suggests that the products, mainly peptides, from the sulfite/sulfate-fueled MSA reaction may have an autocatalytic function. Similar results were observed in the sulfate-fueled MSA reaction treatment groups, with the additional production of cysteine (Figure 6d, e, and f). These results indicate that the peptides exhibit both forward and reverse catalysis. The autocatalysis of peptides generated in MSA reaction network may fit the model A, which is authentic autocatalysis (Scheme 3 a) [39], [40].

**Figure 6.**
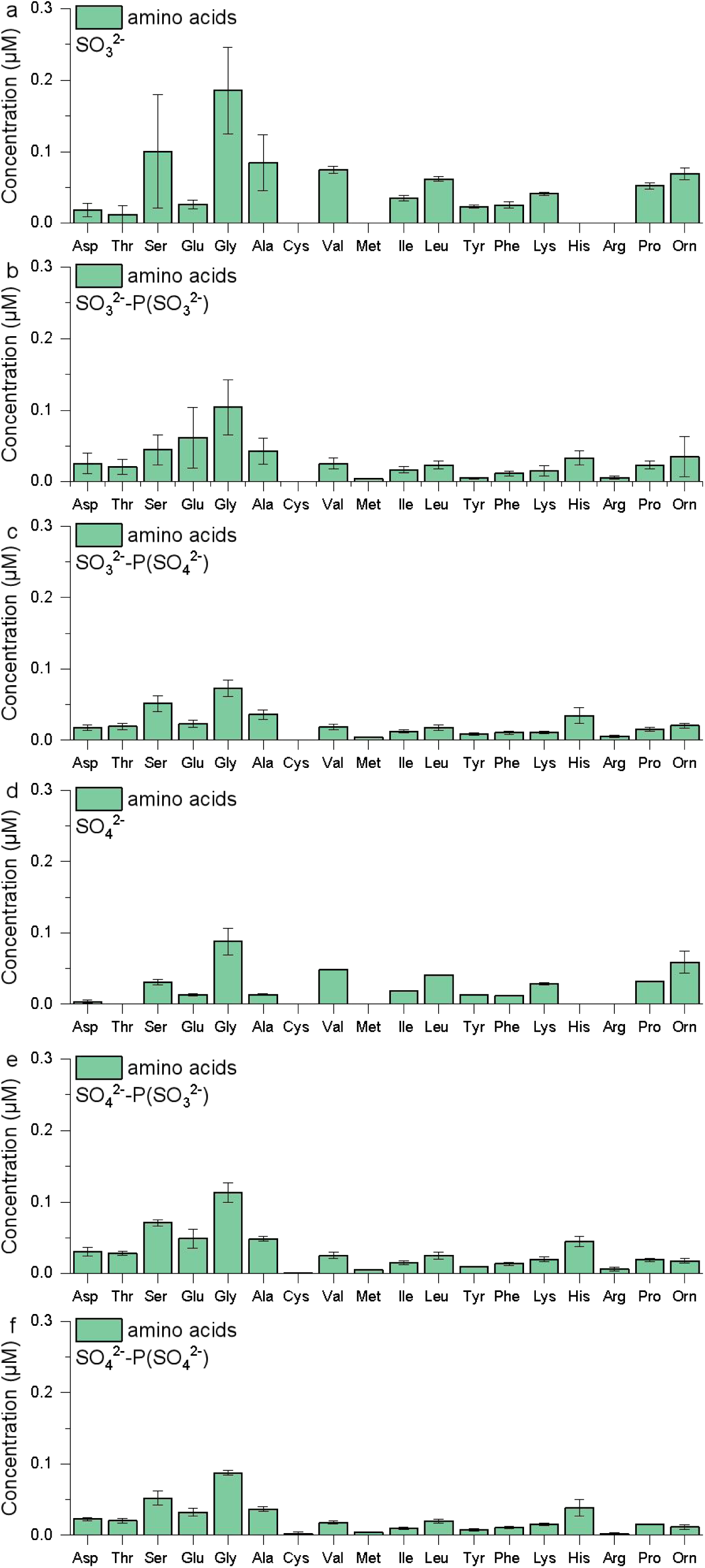
Amino acids generated in MSA reaction network. Treatments were as follows: a, sulfite-fueled MSA reaction network; b, sulfite-fueled MSA reaction network with addition of catalytic products from sulfite-fueled MSA reaction network; c, sulfite-fueled MSA reaction network with addition of catalytic products from sulfate-fueled MSA reaction network; d, sulfate-fueled MSA reaction network; e, sulfate-fueled MSA reaction network with addition of catalytic products from sulfite-fueled MSA reaction network; f, sulfate-fueled MSA reaction network with addition of catalytic products from sulfate-fueled MSA reaction network. Error bars represent standard deviations of three replicates.

**Scheme 3.**
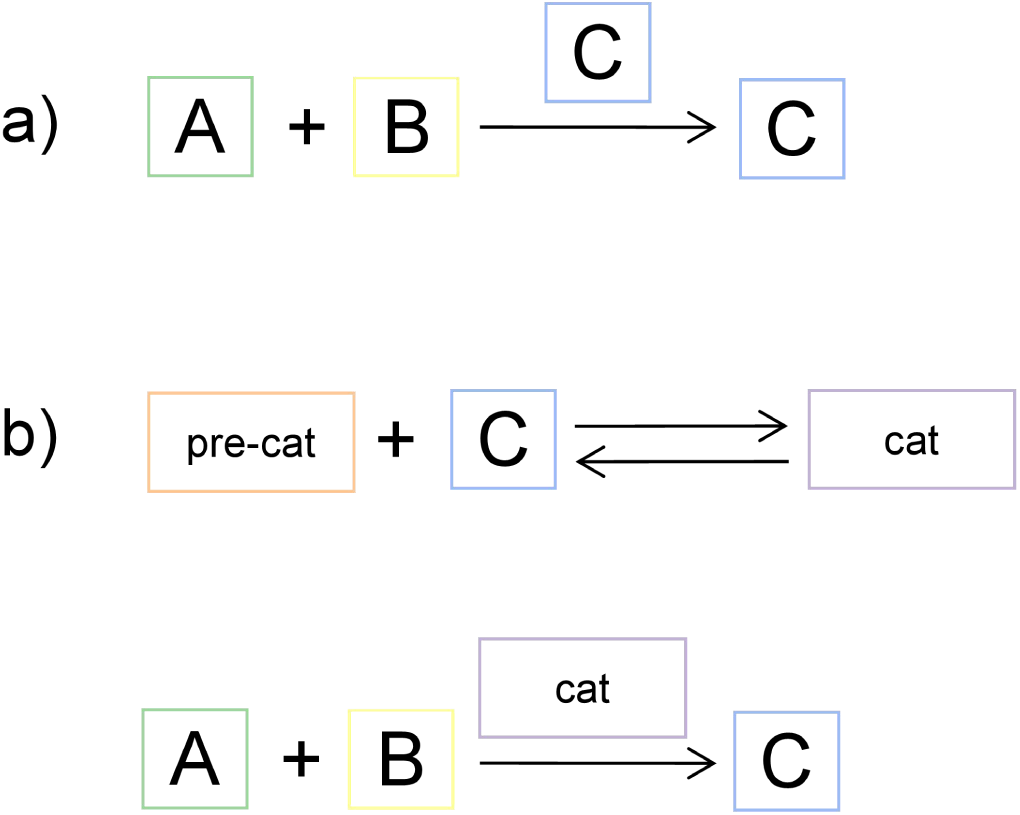
Schematic diagram of autocatalytic reaction mode. (a) Authentic autocatalysis, C (peptides) catalyzes its own formation; (b) catalyst activation, pre-cat (pre-catalyst: Mg^2+^) chelates with C (peptides) to form cat (mature catalyst) over the course of the reaction.

In the experiments outlined in supporting information section E, we conducted a series of investigations to determine if the light-Sammox-driven CRN exhibits autocatalysis. To achieve this, we utilized MSA reaction network products to catalyze the CRN. Specifically, we injected a 1.0 mL reaction solution (the first round light-Sammox-driven CRN with/without Mg^2+^) into a freshly prepared reaction solution (the second round light-Sammox-driven CRN with/without Mg^2+^) as a potential catalyst. The products generated from light-Sammox-driven CRN without Mg^2+^ exhibited tiny reverse catalysis in the treatment groups (Figure 3 and 7a, b). In fact, the addition of Mg^2+^ to light-Sammox-driven CRN resulted in lower peptides production compared to the treatment groups without Mg^2+^ (Figure 7a, b, c). The products generated from light-Sammox-driven CRN with Mg^2+^ treatment groups were found to significantly facilitate the generation of peptides in the second round reaction compared to the first round reaction (Figure 7c and d). Peptides show higher catalytic efficiency through chelation with Mg^2+^ in light-Sammox-driven CRN, and fit the autocatalytic model B (Scheme 3b) [39], [40]. Those results highlighted the critical role of Mg^2+^ in anaerobic photosynthesis origin and evolution in anoxygenic phototrophs [19].

**Figure 7.**
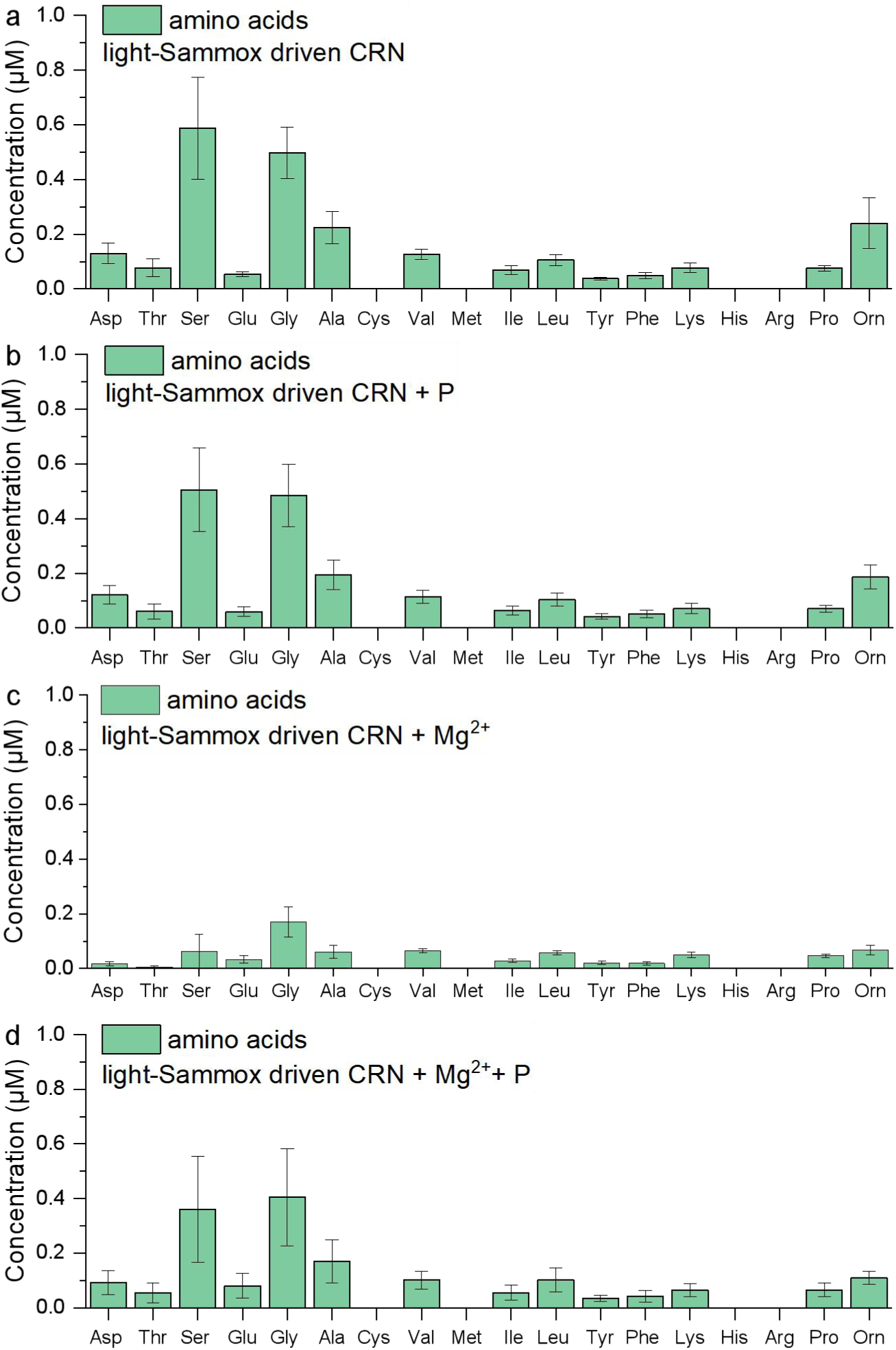
Amino acids generated in light-Sammox-driven CRN. Treatments were as follows: a, light-Sammox-driven CRN; b, light-Sammox-driven CRN with addition of catalytic products from light-Sammox-driven CRN; c, light-Sammox-driven CRN with addition of Mg^2+^; d, light-Sammox-driven CRN with addition of Mg^2+^ + catalytic products from light-Sammox-driven CRN (Mg^2+^ addition treatment groups). Error bars represent standard deviations of three replicates.

**Figure 8.**
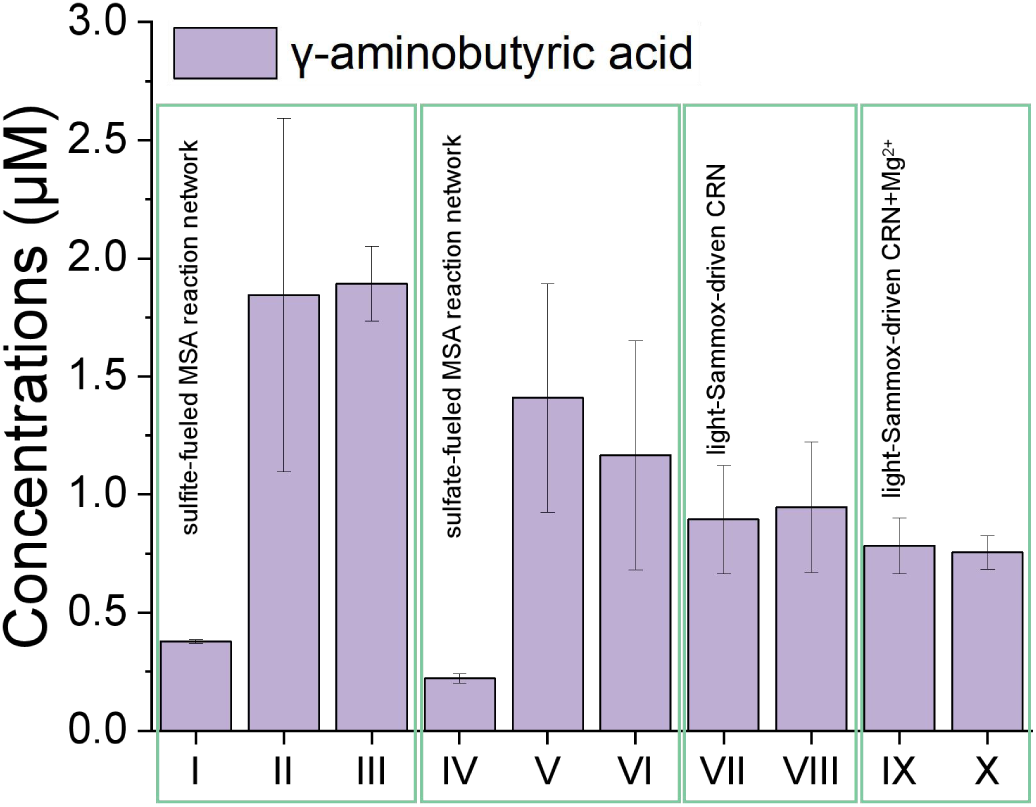
γ-aminobutyric acid generated in MSA reaction network and light-Sammox-driven CRN. I, sulfite-fueled MSA reaction network; II, sulfite-fueled MSA reaction network with addition of catalytic products from sulfite-fueled MSA reaction network; III, sulfite-fueled MSA reaction network with addition of catalytic products from sulfate-fueled MSA reaction network; IV, sulfate-fueled MSA reaction network; V, sulfate-fueled MSA reaction network with addition of catalytic products from sulfite-fueled MSA reaction network; VI, sulfate-fueled MSA reaction network with addition of catalytic products from sulfate-fueled MSA reaction network; VII, light-Sammox-driven CRN; VIII, light-Sammox-driven CRN with addition of catalytic products from light-Sammox-driven CRN; IX, light-Sammox-driven CRN with addition of Mg^2+^; X, light-Sammox-driven CRN with addition of Mg^2+^ + catalytic products from light-Sammox-driven CRN (Mg^2+^ addition treatment groups). Error bars represent standard deviations of three replicates.

#### 2.2.2 weakly reversible realization in MSA reaction network and light-Sammox-driven CRN

We found weakly reversible realization that proteinogenic amino acids (L-serine, glycine, L-alanine, L-valine, L-isoleucine, L-leucine, L-tyrosine, L-phenylalanine, L-lysine, and L-proline) generated in the sulfite-fueled MSA reaction network with catalytic products from the sulfite/sulfate-fueled MSA reaction treatment groups (Figure 6b and c) appear to be slightly less than those in the sulfite-fueled MSA reaction network (Figure 6a). The concentrations of amino acids produced in the sulfite-fueled MSA reaction network, using catalytic products from the sulfite/sulfate-fueled MSA reaction treatment groups (0.49 ± 0.22 and 0.38 ± 0.08 μM), seem to be lower compared to those in the sulfite-fueled MSA reaction network (0.81 ± 0.24 μM) (Table S1). The total concentrations of amino acids generated in the light-Sammox-driven CRN (2.13 ± 0.56 μM) with catalytic products appear to be slightly lower than those in the light-Sammox-driven CRN (2.35 ± 0.61 μM) (Table S1). Except for glutamate and phenylalanine, the concentration of other amino acids in the light-Sammox-driven CRN with catalytic products was all lower than that in the light-Sammox-driven CRN (Figure 7). Similar to Sammox-driven CRNs, the reversibility in the MSA reaction network and light-Sammox-driven CRN may be attributed to the reverse catalysis of peptides.

### 2.3 three main interdependent components of life emerged in MSA reaction network and light-Sammox-driven CRN

#### 2.3.1 the catalytic capacity of energy release reactions in MSA reaction network and light-Sammox-driven CRN

For the MSA reaction network, the products generated during the first round of reactions have been shown to significantly enhance ammonium and methane consumption in all subsequent second-round reactions (Figure 1a and b vs. Figure 5a and b). No significant change was observed in the amount of substrate sulfur, which may be attributed to the relatively rapid cycling and transformation of sulfur within the MSA reaction network. In contrast, for light-Sammox-driven CRN, total substrate sulfur consumption in second-round reactions exceeds that in first-round reactions (Figure 3c). Alternatively, it is possible that new sulfur compounds are being produced. Magnesium ions were found to promote ammonia consumption in light-Sammox-driven CRN (Figure 3a). Similar results were reported in our previous study, indicating that products generated from Sammox-driven CRN can catalyze exergonic chemical reactions such as Sammox and anammox (anaerobic ammonium oxidation coupled with nitrite reduction) [4].

Ornithine, a non-proteinogenic amino acid, was detected in both the MSA reaction network and light-Sammox-driven CRN (Figures 6 and 7). It has been demonstrated to act as a promoter for proto-protein folding and functionalization [41]. The peptides generated from these two CRNs are likely complex catalysts capable of facilitating energy-releasing chemical reactions. Notably, we discovered nitrate generation within Sammox-driven CRN following short-term hydrothermal treatment followed by three years of ambient environmental reaction [42]. As is well known, anaerobic ammonium oxidation to nitrate occurs during microbiological sulfate reduction coupled with anaerobic ammonium oxidation processes. This suggests potential evolutionary developments regarding peptide catalytic functions within Sammox-driven CRN and implies possibilities for generating proto-metabolic machinery.

#### 2.3.2 the potential PNA backbones in MSA reaction network and light-Sammox-driven CRN

The pseudopeptide nucleic acid mimic, known as PNA (peptide nucleic acid), has been proposed as a robust prebiotic evolutionary precursor to RNA. It is capable of facilitating the transfer of chemical sequence information from one PNA oligomer to another—a replicative process—as well as from a PNA oligomer to an RNA oligomer, representing a transition from PNA to RNA [43], [44]. The N-(2-aminoethyl)glycine, ornithine, glycine, and aminobutyric acid are commonly recognized as the foundational backbones of peptide nucleic acids (PNAs) [45–47]. The chiral peptidic DNA composed of glycine and γ-aminobutyric acid demonstrated self-recognition properties akin to those observed in DNA-DNA duplexes [45], [46]. We identified L-ornithine, glycine, and γ-aminobutyric acid in both MSA reaction network and light-Sammox-driven CRN (Figure 6, 7, 8; Figure S2). It is noteworthy that other isomers of ornithine and aminobutyric acid were either not generated or fell below the detection limit. Consequently, the stability of the PNA decamer complexes could be achieved with chirally pure PNA backbones [47]. These results suggest the potential for generating a template/genetic mechanism from the MSA reaction network and light-driven Sammox CRN.

#### 2.3.3 the membrane compartments in MSA reaction network and light-Sammox-driven CRN

Additionally, the stoichiometry approach employed in their study demonstrates the potential for the formation of diverse autocatalytic networks within various compartments, such as porous media, vesicles, and multiphasic systems. We observed the presence of vesicles in the samples obtained from the Light-Sammox-driven CRN and MSA reaction network after 48 hours reaction and 5 months retention (Figure 9). These membranes of vesicles are potentially formed by peptides. While we have not yet confirmed the exact composition, it is believed to provides a compartment to keep its components together and distinguish itself from the environment.

**Figure 9.**
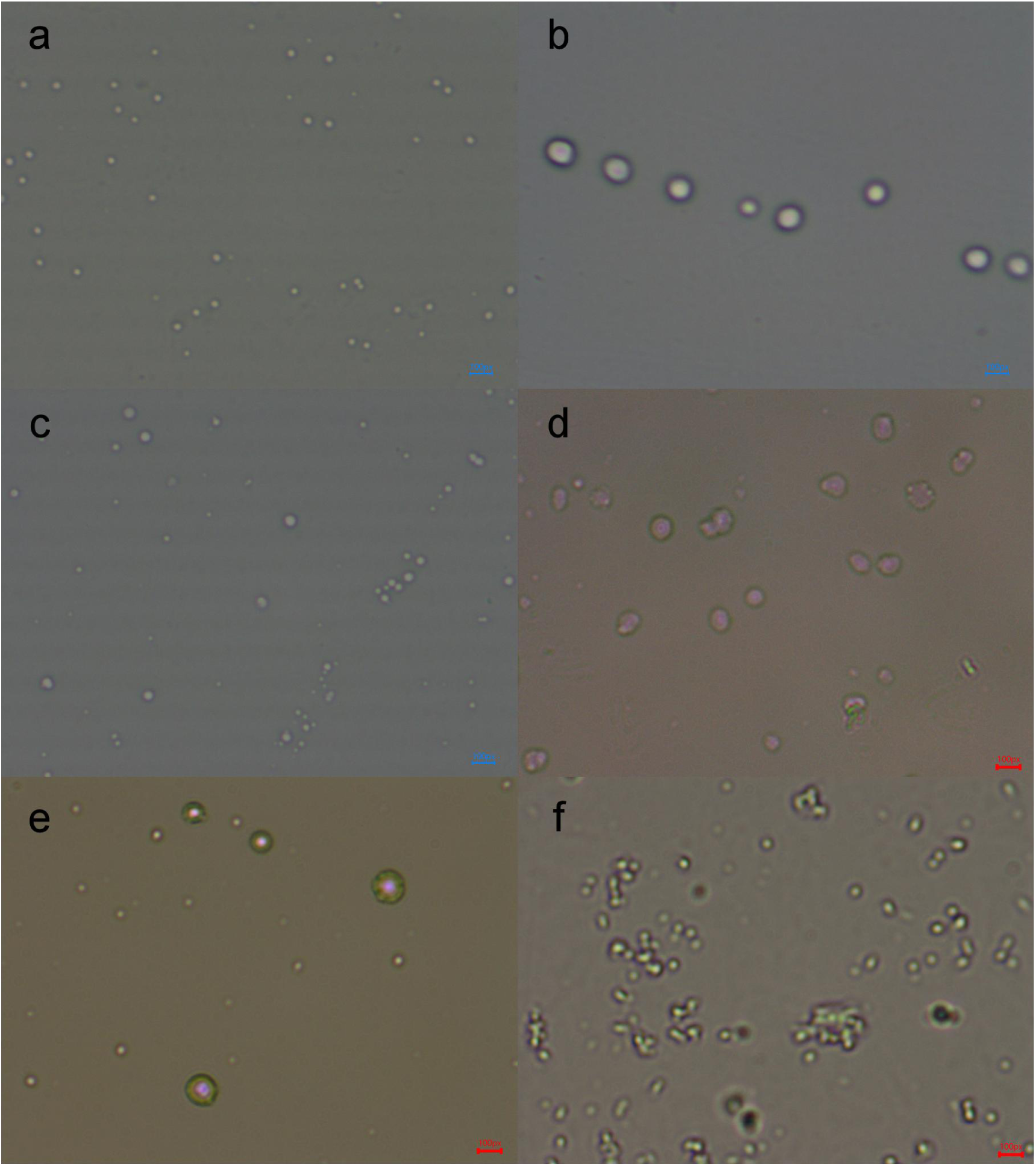
Vesicles formed in sulfite-fueled MSA reaction network after 48 hours reaction (a), in sulfate-fueled MSA reaction network after 48 hours reaction (b), in Light-Sammox-driven CRN after 48 hours reaction (c), Vesicles formed in sulfite-fueled MSA reaction network after 5 months (d). in sulfate-fueled MSA reaction network after 5 months (e), in Light-Sammox-driven CRN after 5 months (f). 100px represents microscope magnification.

### 2.4 the "five nodes" principle as a universal theory of life

#### 2.4.1 theoretical hypothesis and elaboration of the "five nodes" principle in the origin of life

We have demonstrated the potential of NESFBE to give rise to autopoietic systems by identifying autocatalysis, weakly reversible realizations, metabolic machinery, membrane compartments, and template/genetic mechanisms within two CRNs. Our concern is that the "five nodes" principle might closely related with the emergence of autocatalytic networks. Blokhuis et al (2020) found that the minimal motifs of autocatalytic networks, namely, autocatalytic cores, which have five fundamental categories of motifs [30]. These networks exhibit internal catalytic cycles that promote growth. The best example is the reductive TCA cycle, an autocatalysis cycle containing five ‘pillars of anabolism’—acetate (as acetyl-CoA), pyruvate, oxaloacetate, succinate (or succinyl-CoA) and α-ketoglutarate [48]. Our previous study have shown that there were succinate and α-ketoglutarate emerged from Sammox-driven CRN, except for acetate [49]. Furthermore, pyruvate could converted into succinate, α-ketoglutarate, and proteinogenic amino acids suggesting an autocatalysis proto-rTCA reaction network in Sammox-driven CRN [49]. In fact, pyruvate, oxaloacetate, and α-ketoglutarate could all reacted with the product of Sammox-driven CRN—ammonium formate to generate up to 13 kinds of proteinogenic amino acids [4]. There might be five ‘pillars of anabolism’ generated in Sammox-driven CRN as intermediates during the synthesis of amino acids from carbon dioxide, although we haven’t detected pyruvate and oxaloacetate directly in Sammox-driven CRN.

There could be a potential hierarchical organization in the evolution of complex networks during the origin of life [50]. Scheme 4 presents a conceptual model illustrating the "five nodes" principle of this hierarchical network, ranging from the inorganic molecular level to the motif level, ultimately leading to autocatalysis. Scheme 4a represents various CRNs, such as light-Sammox-driven CRN and MSA reaction network, all of which exhibit a highly integrated connection between C, H, O, N, and S. Scheme 4b, on the other hand, represents an organic compounds reaction network that is generated hierarchically from Scheme 4a. These replication and connection steps can be repeated indefinitely, with each step increasing the number of nodes in the system by a factor of 5 [51]. In fact, there is the "five nodes" principle at a higher level than motif level, for instance the *Escherichia coli* metabolic network contains five highly connected nodes: glutamate, coenzyme A, 2-oxoglutarate, pyruvate, and glutamine [52].

**Scheme 4.**
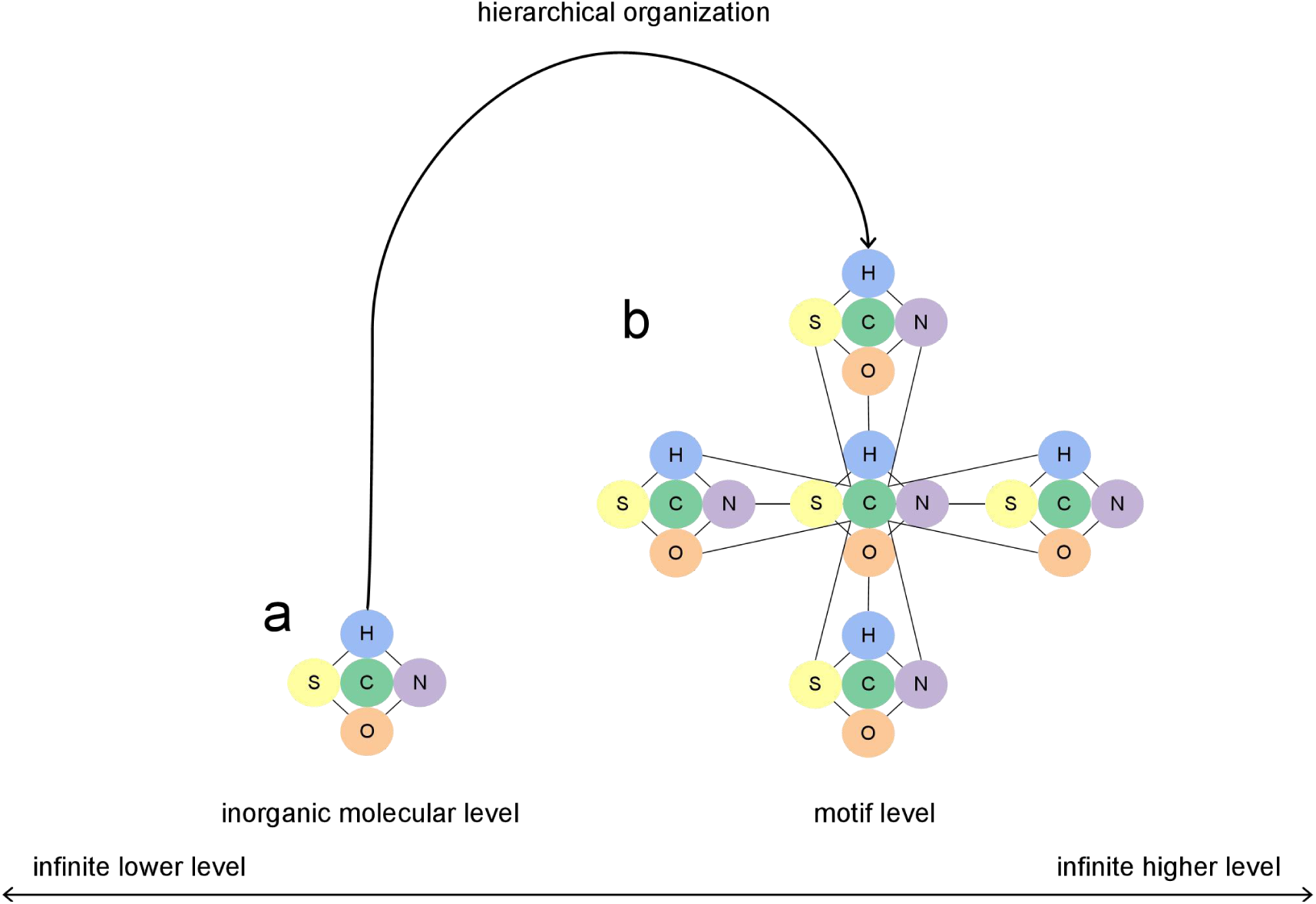
The conceptual model of the "five nodes" principle in hierarchical network leading to autocatalysis. The "five nodes" principle at the inorganic molecular level are represented by highly integrated five-node modules(a). At the motif level, there are 25-node modules representing the "five nodes" principle (b). To create a network of five super-nodes (motifs), four identical replicas connect the peripheral nodes of each cluster to the central node of the original cluster. Not all nodes will be displayed due to overlap and occlusion. Auxiliary nodes/elements are not shown. 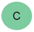, carbon; 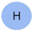, hydrogen; 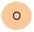, oxygen; 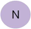, nitrogen; 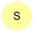, sulfur.

Hierarchy is a fundamental characteristic of many heteropoietic systems, such as the Worldwideweb, actor network, the Internet at the domain level, and the semantic web [51], [53]. In this study, we realized that the mathematical principles of hierarchically repeated five nodes in autopoietic systems probably came from the Sharkovsky’s cycle coexistence ordering. Let *F* be a continuous function from the real interval *J* to *J* itself. For any positive integers *m* and *n*, if *m* comes after *n* in the Sharkovsky’s cycle coexistence ordering, then the continuous function *F*: *J*→*J* must have periodic points of period *n* as long as it has periodic points of period *m* [54].

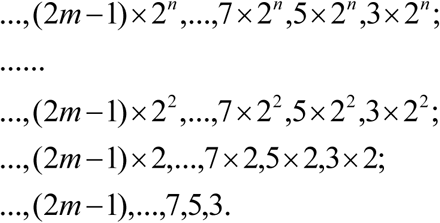

As widely acknowledged, Li and Yorke (1975) established that period three signifies chaos [55]. Chaos is an inherent characteristic of autocatalytic systems [56]. It logically follows that periodic points of period three must manifest at the inception of life. Consequently, we can infer the pervasive existence of period five throughout the entire timeline of living systems. The presence of period five indicates autopoiesis, extending from autopoietic to heteropoietic systems. The Sharkovsky’s cycle coexistence ordering is the mathematical basis for the inevitable emergence of life.

The genesis of the numeral "5" in the "five nodes" principle, can be comprehended through the application of graph theory. This is because modern thermodynamics of discrete systems relies on graph theory, which offers algebraic methods to define observables and a geometric intuition of their meaning and role. Dal Cengio et al (2023) discovered that by directly imposing Kirchhoff’s rules and linearizing the system, the thermodynamic feasibility of reconstructing metabolic networks can be achieved [57]. Consequently, attaining a feasible state in the reconstruction of metabolic networks is equivalent to finding boundary solutions within the linear regime. This implies that the number of spanning trees should be five [57]. We deduce that the number five in this context is derived from the linear response of the five degrees of freedom of an ellipse. An ellipse can be fully described by five constraints, making it the maximum number of constraints for which a minimal ellipse can still be sought. When five points are used, a conic can be fully described with no degree of freedom left for area minimization. Equation (8) is commonly used to represent an ellipse, and it provides four degrees of freedom, namely the semi-major axis (*a*), semi-minor axis (*b*), and the center coordinates (*x*_0_, *y*_0_) of the ellipse. However, equation (8) only represents an ellipse with its major and minor axes parallel to the coordinate axes.

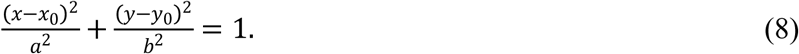

An ellipse with a specific orientation can only be defined by five degrees of freedom: the semi-major axis (*a*), semi-minor axis (*b*), center coordinates (*x*_0_, *y*_0_), and the direction angle (*θ*) [58], [59]. Essentially, the problem arises from the five degrees of freedom associated with the (normalized) coefficients of the ellipse [60].

In our opinion, the five minimal motifs, which can be observed as connected subgraphs in Scheme 4b, consist of five highly integrated nodes. This indicates that each minimal motif contains no more than five kinds of elements, a character that is fulfilled in Sammox-driven CRN and MSA reaction network. Considering that autocatalysis is the foundation of metabolic networks, the significance of the number five in the generation of autocatalysis is revealed.

Let us further explore the "five nodes" principle at the inorganic molecular level (Scheme 4a). Similarly, a CRN can be represented by an oriented linear graph [61], [62]. For instance, when considering a directed linear graph *g*, the cutset matrix *Q* (Eq. 9) and loop matrix *B* (Eq. 10) can be defined as follows:

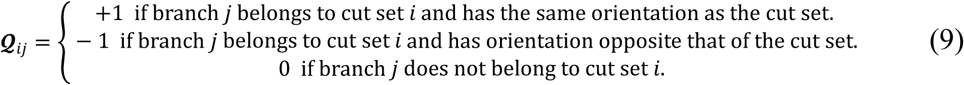

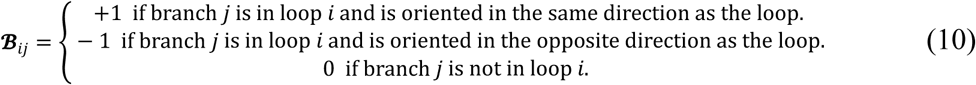

The computation of dynamically equivalent detailed balanced and complex balanced CRNs can be traced back to simple linear programming [63]. Therefore, the "five nodes" principle at the inorganic molecular level can also be derived from the normalized coefficients of an ellipse, as proposed by [60], [64].

The requirement of weak reversibility in CRNs can be formulated as a linear constraint within the framework of mixed-integer linear programming [31]. Weak reversibility implies that every reaction in the network can be reversed by a suitable sequence of subsequent reactions. It is important to note that not every reaction needs to have a reverse reaction for the system to be weakly reversible, as illustrated by the concept of weakly reversible networks introduced by Johnston and Siegel (2011) [65]

**Scheme 5.**
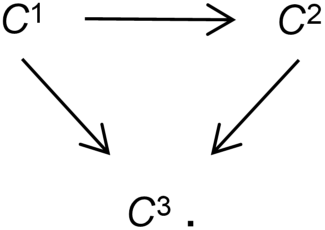
The conceptual model of the weakly reversible network [65].

#### 2.4.2 the weakly reversible realization upgraded the concept of autopoiesis

In a previous study, it was discovered that the peptides produced in Sammox-driven CRNs demonstrate a unique phenomenon of exhibiting both forward and reverse catalysis [4]. Interestingly, this catalytic behavior differs depending on whether the CRNs are fueled by sulfite or sulfate. Furthermore, this distinction in catalytic impact is observed across both a variable temperature range and a fixed temperature, ultimately leading to the emergence of seesaw-like catalytic properties. These findings shed light on the intricate dynamics and complexities of Sammox-driven CRNs. The redox property of Sammox-driven CRNs can be influenced by variations in substrate concentration and temperature. Such changes have the potential to disrupt the redox homeostasis of the system. Consequently, the catalytic orientation of peptides undergoes alterations in response to these stimuli, ensuring the maintenance of system redox homeostasis. This seesaw-like catalytic behavior of peptides plays a crucial role in achieving the sustainability and evolution of Sammox-driven CRNs [4], [5]. Moreover, the reverse catalytic properties of peptides offer a valuable source of reversibility for Sammox-driven CRNs. In this study, we observed weakly reversible realizations within the Sammox-driven CRN and MSA reaction networks. Those results support a linear constraint on CRNs concerning the origin of life, which serves as the foundation for the "five nodes" principle that governs this process.

Autocatalytic chemical reaction systems can be regarded as reaction-diffusion systems involving both activators and inhibitors. An activator performs two key functions: first, it promotes its own production (autocatalysis); second, it induces the formation of its corresponding inhibitors. Inhibitors are generated in response to the activator but act to suppress the activator’s activity [66], [67]. In our study, activators may correspond to autocatalytic peptides, while inhibitors could be peptides exhibiting reverse catalytic activity. Since chaos and fractals are inherent characteristics of reaction-diffusion processes [68], our weakly reversible realization CRNs exhibits chaotic behavior, specifically manifesting as period-three dynamics. This further provides a foundation for the emergence of period-five.

The integration of data derived from our chemical studies and mathematical physical derivations provides compelling evidence for the mathematical principles underlying the origin of life. This principle suggests that levels of replication and connection can be perpetuated indefinitely within hierarchical autopoietic systems (Schemes 4 and 6). Consequently, all autopoietic systems may be regarded as components of the larger system they collectively constitute (Scheme 6). The maintenance of an autopoietic system necessitates that its components are continuously or periodically produced and disintegrated [69]. The disintegration — such as biomolecular degradation, cell apoptosis, tissue necrosis, and individual death — of a component within an autopoietic system signifies the demise of one or more subordinate autopoietic systems (Scheme 6). The essence of disintegration is characterized by weakly reversible realizations (Scheme 6), which are crucial for both the formation and maintenance of autopoietic systems. We propose that autopoietic systems should be defined by their capacity for self-production, self-maintenance, and self-disintegration. Thus, we infer that linear constraints govern both the emergence of the autopoietic system (life) and its eventual self-disintegration (death) in a hierarchical manner. The "five nodes" principle elucidates the inevitability with which life confronts death, emphasizing that death is an intrinsic aspect of life; without death, life cannot be fully realized.

**Scheme 6.**
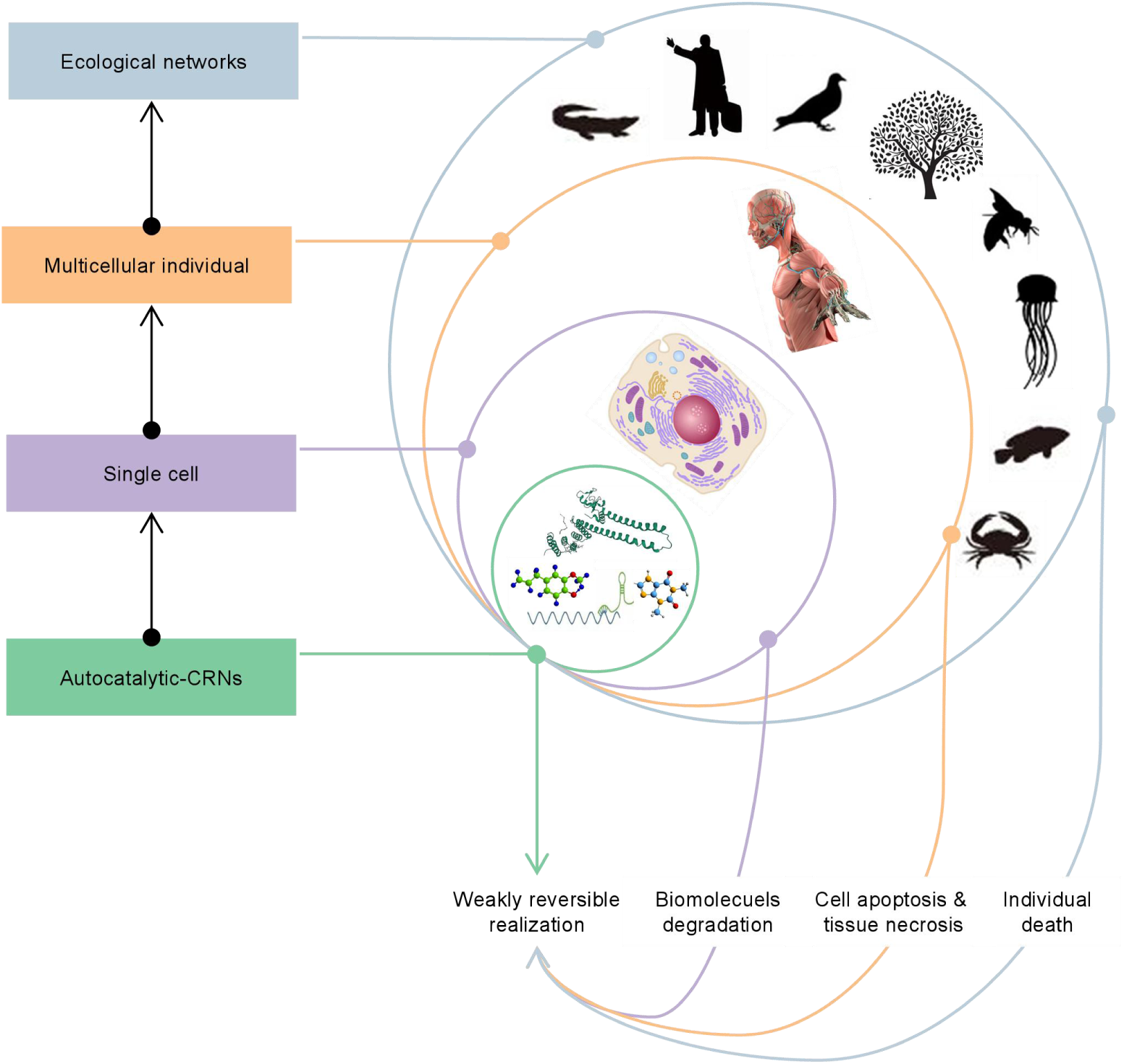
Hierarchical self-assembly of autopoietic systems, encompassing autocatalytic CRNs to ecological networks, and the manifestation of weakly reversible realizations (disintegration), ranging from biomolecules degradation to individual death at various levels within the autopoietic system.

## 3. Conclusions

We posited that the non-equilibrium synergy of five bioessential elements (NESFBE) could play a significant role in the emergence of life. There may exists a "five nodes" principle as a universal theory of life, which may closely associated with the emergence and progression of autopoietic systems. In this study, we report on the generation of autocatalytic peptides from both the MSA reaction network and light-Sammox-driven CRN. Furthermore, products (mainly peptides) derived from the MSA reaction network and the Light-Sammox-driven CRN exhibit a distinctive reverse catalytic capability. This characteristic provides weakly reversible realizations for each CRN— a feature crucial for imposing linear constraints on these networks. We also found the possible emerge of the three main interdependent components of life in the Light-Sammox-driven CRN and MSA reaction network. The peptides generated from the two CRNs could catalysts the energy-releasing chemical reactions and facilitate substrates consumption. L-ornithine, glycine, and γ-aminobutyric acid are as the common PNA backbones in both MSA reaction network and light-Sammox-driven CRN, implying the possibility of generating template/genetic mechanism. The vesicles formed in the Light-Sammox-driven CRN and MSA reaction network have the potential to facilitate autocatalysis. The NESFBE-based chemical reaction networks, comprises NH_4_^+^, SO_3_^2-^/SO_4_^2-^, HCO_3_^-^, CH_4_, and H_2_ O, establishes an intricate network conducive to the origin of methanotrophy and anoxygenic phototrophy.

Our findings provide evidences supporting a linear constraint on chemical reaction networks that is pivotal to the origin of life. The number five in the "five nodes" principle arises from this linear constraint. The "five nodes" principle posits the potential for indefinite replication and connection steps within hierarchically organized living systems, where each step increases the number of nodes in the system by a factor of five. In this context, hierarchical organization within living systems corresponds to mathematical periods. In conclusion, “period five indicates autopoiesis” could be a universal theory of life. Linear constraints guiding both the emergence and demise of the autopoietic system.

## METHODS

Detailed methods can be found in the supplemental information.

## SUPPLEMENTAL INFORMATION

Supplemental information can be found online.

## AUTHOR CONTRIBUTIONS

Peng Bao conceived the study, designed and carried out the experiment, and wrote the manuscript. Min Qiu carried out experiments and analysis.

## DECLARATIONS

### Ethics approval and consen to participate

Not applicable.

### Consent for publication

Not applicable.

