## supplemental table and figures for "Period five indicates autopoiesis"

***Corresponding Author**

**Supporting information**

**MATERIAL AND METHODS**

**A. Chemicals and reagents**

All chemical reagents and organic solvents used in this study were of analytical grade. Ammonium chloride (>99.5%, CAS number: 12125–02–9) was obtained from Sigma–Aldrich, USA, while sodium sulfate (>99.9%, CAS number: 7757-82-6) was purchased from Aladdin, USA. Sodium sulfite (>98.5%, CAS number: 7757-83-7) was purchased from Acros Organics, Belgium. Sodium formate, (99.5%, CAS 141-53-7) was purchased from Rhawn, China. Sodium acetate anhydrate (99.0%, CAS 127-09-3) was purchased from Aladdin, USA. Sodium bicarbonate (>99.5%, CAS number: 144-55-8) was purchased from Sinopharm, China. Methanol (>99.9%, CAS number: 67–56–1) was obtained from Tedia, USA. The purity of methane is 99.9%. Unless otherwise noted, all reagents were used without further purification, and ultrapure water was prepared using the Millipore purification system. (Billerica, MA, USA).

**B. General procedure for MSA reaction network**

One hundred milliliters of ultrapure water were added to 120 mL serum bottles and sealed with butyl rubber stoppers and aluminum crimp caps. The solution was then autoclaved and cooled to 25°C after being flushed with 99.999% pure helium gas. To create the "MSA reaction system," additional sulfite/sulfate, ammonium, and methane were aseptically introduced to the serum bottles. Sulfite/sulfate (1 mL, 3 mM final concentration), ammonium solution (0.5 mL, 4 mM final concentration), and methane (1 mL, 1 mM final concentration) were added to the serum bottles. The initial pH value is about 7.7. The reaction systems were then heated at 70°C in a water bath for 48 hours in the dark. After this, the systems were removed from the water bath and allowed to cool to room temperature before the investigation was conducted. The investigation involved a series of experiments, which are detailed below:

(i) 3 mM sulfate + 4 mM NH_4_Cl + 1 mM CH_4_,
(ii) 3 mM sulfite + 4 mM NH_4_Cl + 1 mM CH_4_,

**C. General procedure for light-Sammox-driven CRN and Sammox-driven CRN**

**1. General procedure for light-Sammox-driven CRN**

To prepare the "light-Sammox-driven CRN," 100 mL of ultrapure water was added to 120 mL serum bottles, which were then sealed with butyl rubber stoppers and aluminum crimp caps. The solution was autoclaved and cooled to 25°C after being flushed with 99.999% pure helium gas. Sulfite, ammonium, and bicarbonate were added to the serum bottles in aseptic conditions. Specifically, 1 mL of sulfite was added to achieve a final concentration of 3 mM, 0.5 mL of ammonium solution was added to achieve a final concentration of 10 mM, and 1 mL of bicarbonate was added to achieve a final concentration of 20 mM. The initial pH value is about 8.2. The reaction systems were then subjected to heating at 70°C in a water bath, under fluorescent light with a wavelength range of 400 nm-700 nm, for a duration of 135 hours. Subsequently, the systems were allowed to cool down to room temperature before conducting the investigation. The investigation was carried out using the following experimental setup:

(i) 3 mM sulfite + 10 mM NH_4_Cl + 20 mM HCO_3_^-^ + light

**2. The distinction between light-Sammox-driven CRN and Sammox-driven CRN**

To differentiate between the two CRNs, we evaluated the efficiency of proteinogenic amino acids synthesis by each and conducted the experiment at a lower temperature to minimize the impact of the hydrothermal reaction. The light-Sammox-driven CRN reaction system was then subjected to heating at 30°C in a water bath, under fluorescent light with a wavelength range of 400 nm-700 nm, for a duration of 135 hours. The preparation procedure for Sammox-driven CRN is the same as for light-Sammox-driven CRN. The Sammox-driven CRN reaction system was then subjected to heating at 30°C in a water bath in the dark for a duration of 135 hours. Subsequently, the systems were allowed to cool down to room temperature before conducting amino acids determination. The investigation was carried out using the following experimental setup:

(i) 3 mM sulfite + 10 mM NH_4_Cl + 20 mM HCO_3_^-^ + light
(ii) 3 mM sulfite + 10 mM NH_4_Cl + 20 mM HCO_3_^-^

**D. Verifying of autocatalysis and weakly reversible realizations in MSA reaction network**

The experimental procedure was conducted in two rounds. In the first round, MSA reaction solutions were prepared in serum bottles and heated at 70°C in a dark water bath for 48 hours. The solutions were then left to cool to room temperature. From the first round, one milliliter of the sulfite/sulfate-fueled MSA reaction solution was extracted and injected into newly prepared MSA reaction solutions (the second round) as a potential catalyst. The second round of the Sammox reaction was executed at 70°C for 48 hours and allowed to cool to room temperature before sampling. The experimental process is illustrated in the diagram below. (Scheme S1).

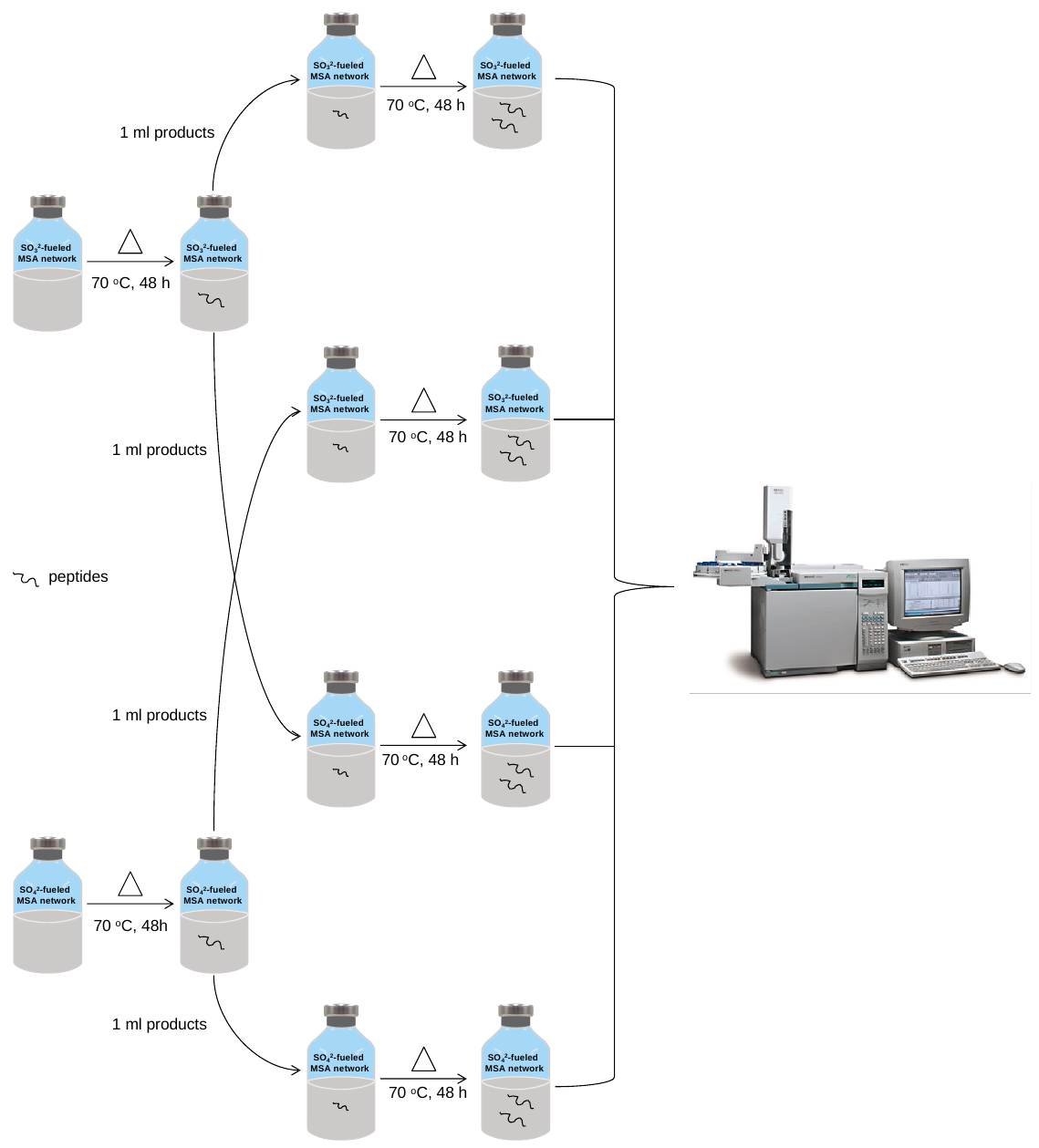

**Scheme S1. Diagram of experimental procedure for verifying of autocatalysis in MSA reaction network.**

**E. Verifying of autocatalysis and weakly reversible realizations in light-Sammox-driven CRN**

The experimental procedure was conducted in two rounds. In the first round, serum bottles containing substrates for the Sammox-driven CRN reaction were heated at 70°C on an electric heating plate under fluorescent light with a wavelength range of 400 nm to 700 nm for 135 hours. The bottles were then left to cool to room temperature. From the first round reaction solutions, one milliliter was extracted and injected into newly prepared Sammox-driven CRN reaction solutions for the second round, serving as a potential catalyst. The second round Sammox reaction was executed in the same manner as the first round reaction and allowed to cool to room temperature before sampling. The experimental process is illustrated in Scheme S2 below.

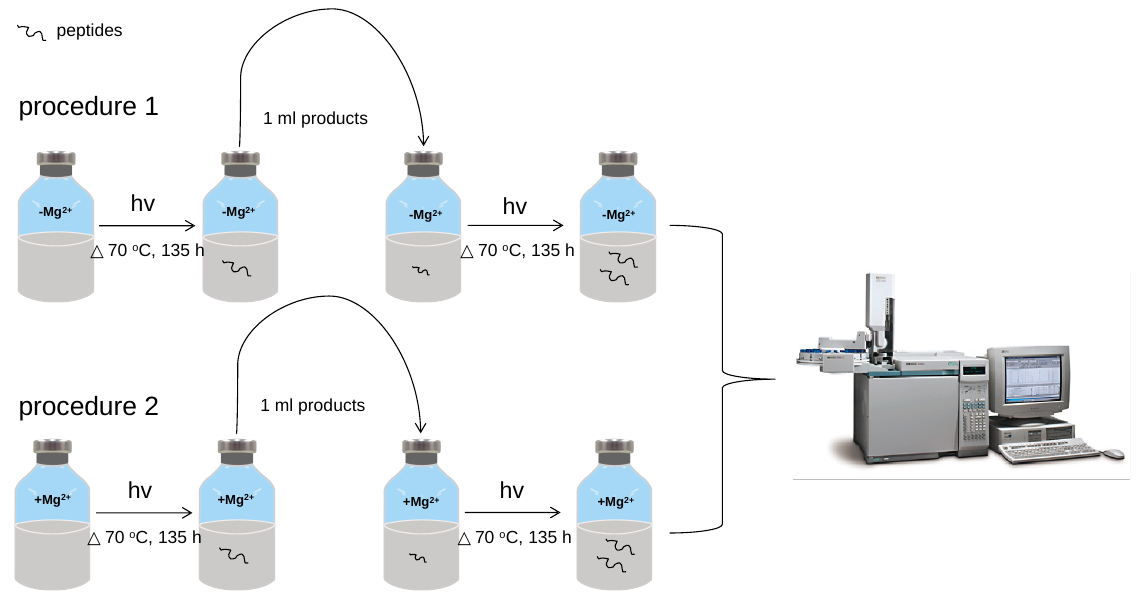

**Scheme S2. Diagram of experimental procedure for verifying of autocatalysis in light-Sammox-driven CRN. Procedure 1, light-Sammox-driven CRN without Mg^2+^. Procedure 2, light-Sammox-driven CRN with Mg^2+^.**

F. **Investigation of membrane compartments, peptide nucleic acids (PNA) backbones, and catalytic capacities for energy release reactions in light-Sammox-driven CRN and MSA reaction network**

The specific details of the investigation of membrane compartments, peptide nucleic acids (PNA) backbones, and catalytic capacities for energy release reactions are shown in sampling analytical methods.

G. **Sampling analytical methods**

**1. Nano LC-MS/MS identification of peptides and amino acids**

Solution samples were freeze-dried and diluted with ultrapure water to 1.0 mL. The sample solution was then reduced using 10 mM DTT at 56°C for 1 hour and alkylated with 20 mM IAA at room temperature in the dark for 1 hour. The extracted peptides were subsequently lyophilized to dryness and resuspended with 2–20 μL of 0.1% formic acid before LC-MS/MS determination. For more detailed information on the method, please refer to the document study by Bao et al. (2022) [1].

For the analysis of amino acids (L-aspartate, L-threonine, L-serine, L-glutamate, glycine, L-alanine, L-cysteine, L-valine, L-methionine, L-isoleucine, L-leucine, L-tyrosine, L-phenylalanine, L-lysine, L-histidine, L-arginine, L-proline, L-ornithine, and γ-aminobutyric acid ), samples solution were freeze-dried and diluted with ultrapure water to 1.0 ml, followed by acid hydrolysis. The detailed analytical method is outlined in the document study by Bao et al. (2022) [1].

**2. Ion chromatography quantitative analysis of sulfite, sulfate, and ammonium.**

The ion chromatography system used in this study consisted of an ICS-5000+ SP pump (Thermo Fisher Scientific Inc., Sunnyvale, CA, USA), a column oven ICS-5000 DC, and an electrochemical detector DC-5. To detect sulfite and sulfate, a Dionex Ionpac AS11-HC column was employed, with a flow rate of 1.0 mL min^-1^ and an eluent of 30 mM KOH. For the detection of ammonium, a Dionex IonpacTM CS 12A column was utilized, with a flow rate of 1.0 mL min^-1^ and an eluent of 20 mM sulphonethane.

**3. Quantitative analysis of thiosulfate by high performance liquid chromatography**

Thiosulfate was quantified using high performance liquid chromatography (HPLC) with a quaternary pump (Agilent 1260 infinity HPLC system, USA). The detailed methodology is outlined in the study document by Li et al. (2020) [2].

**4. Gas chromatography quantitative analysis of methane**

The gas in the reaction bottle was purged with nitrogen using an evacuation cleaning device. Methane was then added to the bottle using a syringe, resulting in an initial methane concentration of 0.955 mmoL L^-1^. The methane concentration in the headspace of the serum bottle was determined using an Agilent 7890A greenhouse gas chromatography system (Palo Alto, CA, USA) by injecting 100 μL samples. The system was equipped with both a flame ionization detector (FID) and a thermal conductivity detector (TCD). The FID parameters were set to a temperature of 250 °C, with an H_2_ flow rate of 35 mL min^-1^, an air flow rate of 395 mL min^-1^, and a He flow rate of 5 mL min^-1^. The TCD parameters were set to a temperature of 200 °C, with an air flow rate of 15 mL min^-1^ and a He flow rate of 1 mL min^-1^. The injection port and column oven temperatures were set to 250 and 60 °C, respectively.

5. **Observation of the vesicles formed in MSA reaction network and light-Sammox-driven CRN by optical microscope**

To identify membrane compartments within the MSA reaction network and the light-driven Sammox CRN. The liquid samples were extracted from the serum bottles after 48 hours reaction. The sample was then kept at 35^o^C in the dark for five months before being sampled again. The sample is diluted with sterile water in equal proportions to remove possible salt crystals. Subsequently, the samples were examined under a Nikon Eclipse E100 microscope at a magnification of ×100, and the resulting images were captured using the ImageView software.

**Table S1. The total concentrations of amino acids generated in light-Sammox-driven CRN and MSA reaction network, and are in accord with that in figure 6 and 7.**

| CRNs | Treatments | Concentrations (μM) |
| --- | --- | --- |
| light-Sammox-driven CRN | light-Sammox-driven CRN | 2.35 ± 0.61 |
|  | light-Sammox-driven CRN+P | 2.13 ± 0.56 |
|  | light-Sammox-driven CRN+Mg^2+^ | 0.70 ± 0.24 |
|  | light-Sammox-driven CRN+Mg^2+^+P | 1.73 ± 0.79 |
| MSA reaction network | SO_3_^2-^ | 0.81 ± 0.24 |
|  | SO_3_^2-^-P(SO_3_^2-^) | 0.49 ± 0.22 |
|  | SO_3_^2-^-P(SO_4_^2-^) | 0.38 ± 0.08 |
|  | SO_4_^2-^ | 0.40 ± 0.05 |
|  | SO_4_^2-^-P(SO_3_^2-^) | 0.54 ± 0.08 |
|  | SO_4_^2-^-P(SO_4_^2-^) | 0.41 ± 0.06 |

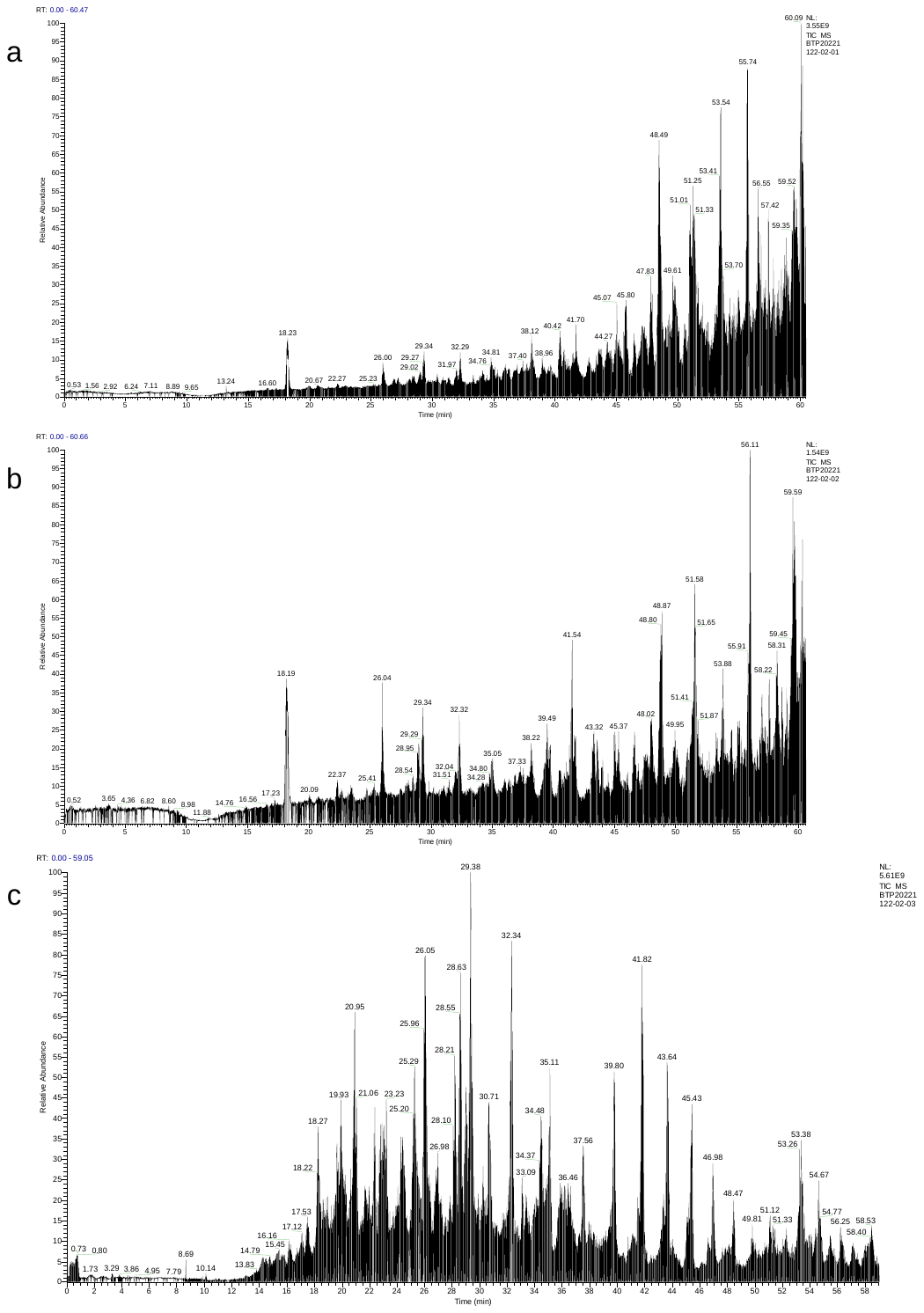

**Figure S1. Total ion flow chromatography of peptides in MSA reaction network and light-Sammox-driven CRN. (a), total ion flow chromatography of peptides generated from sulfite-fueled MSA reaction network; (b), total ion flow chromatography of peptides generated from sulfate-fueled MSA reaction network; (c), total ion flow chromatography of peptides generated from light-Sammox-driven CRN.**

| a | 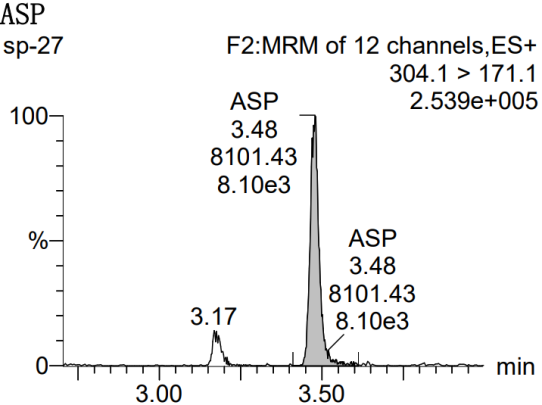 | b | 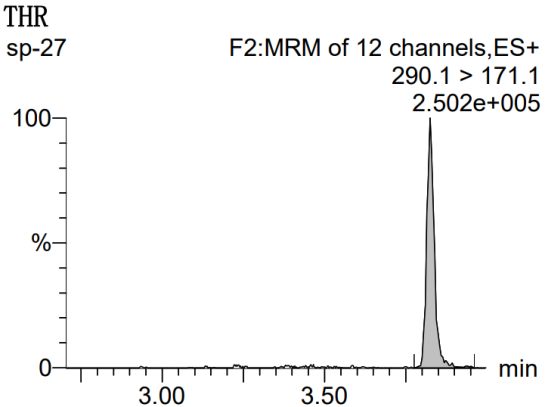 |
| --- | --- | --- | --- |
| c | 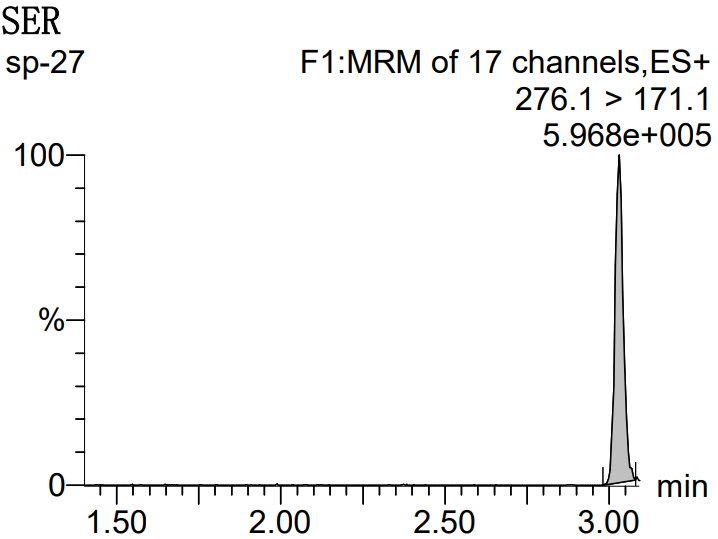 | d | 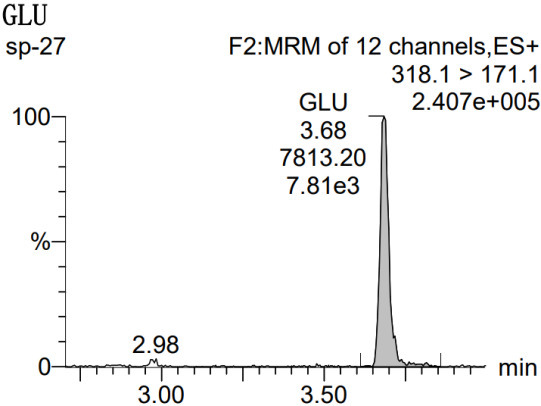 |
| e | 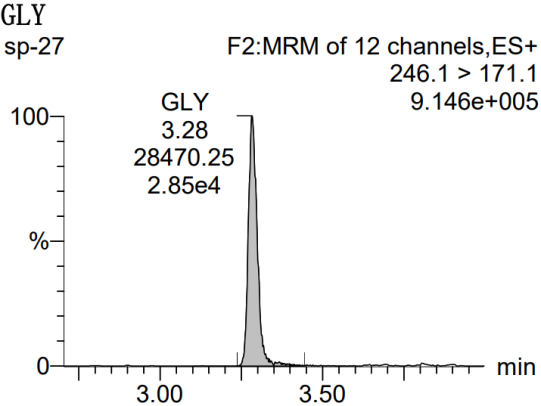 | f | 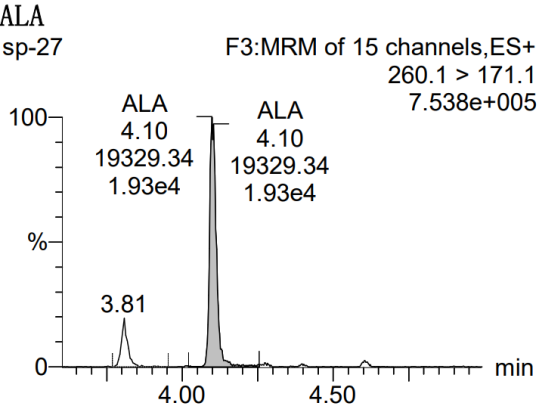 |
| g | 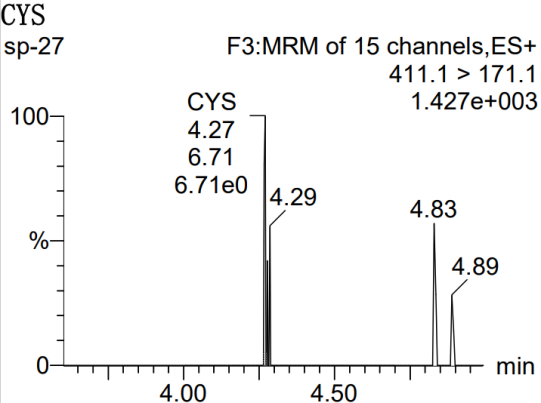 | h | 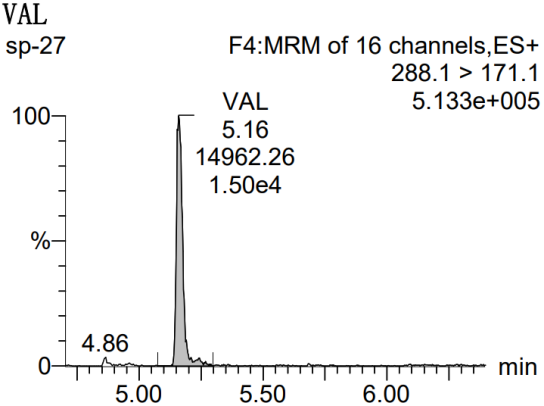 |
| i | 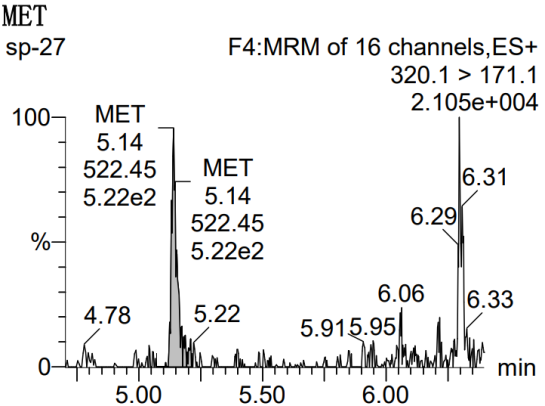 | j | 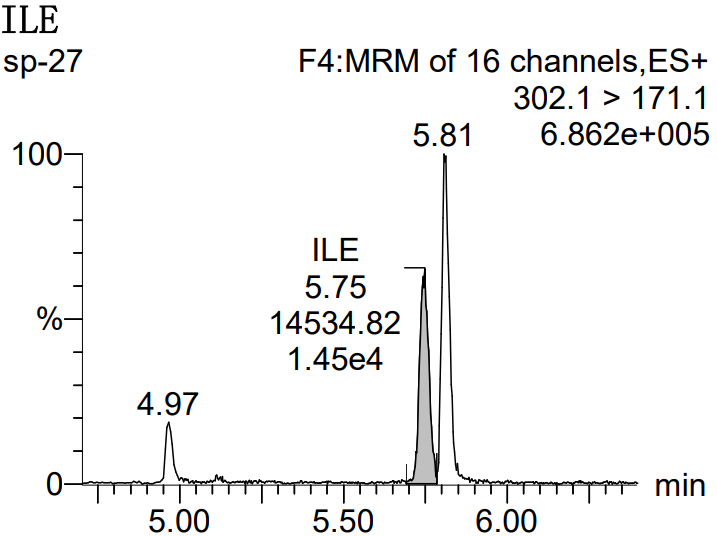 |
| k | 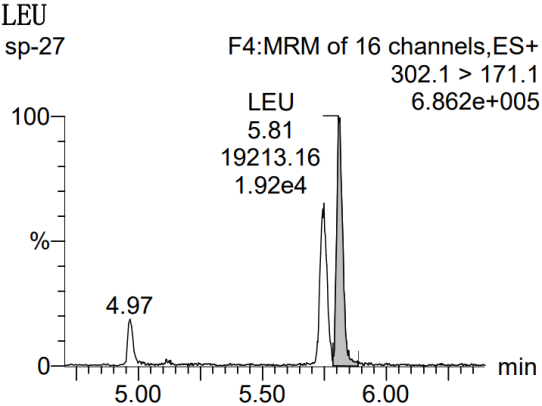 | l | 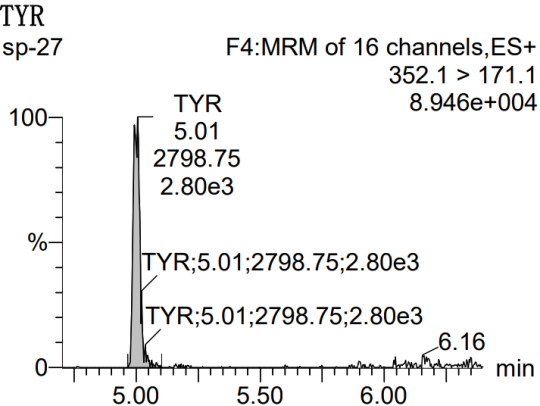 |
| m | 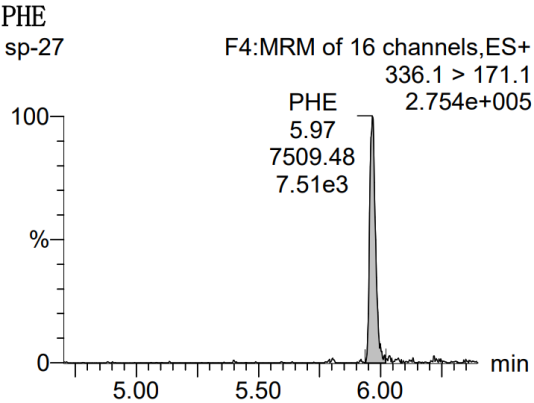 | n | 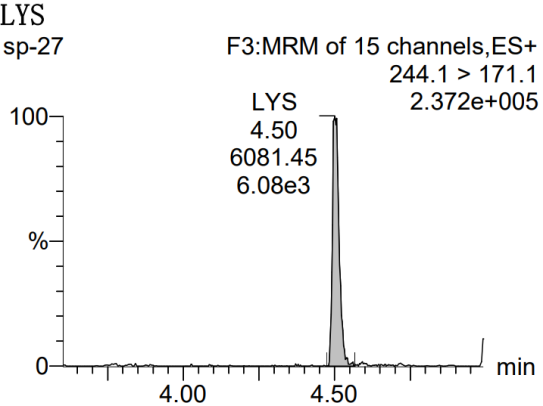 |
| o | 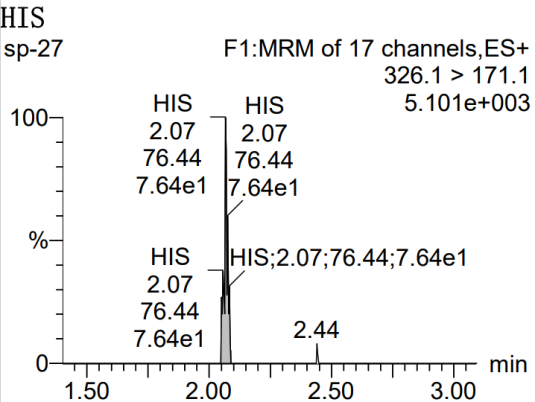 | p | 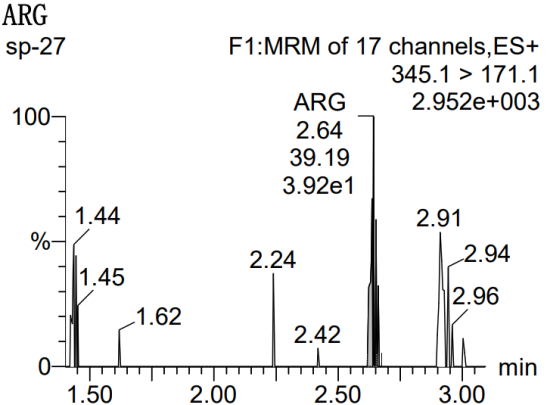 |
| q | 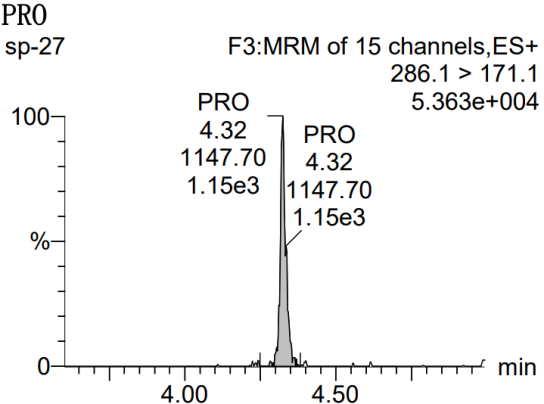 | r | 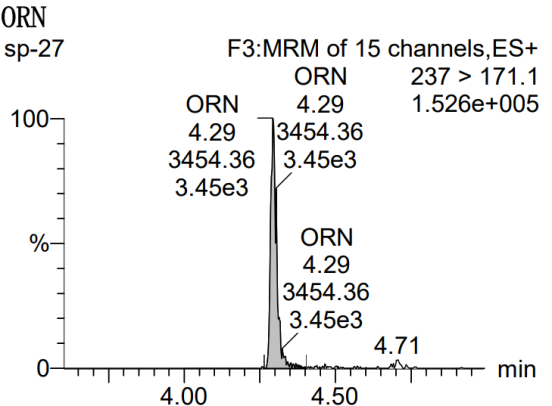 |

**Figure S2. Chromatogram of amino acids detected in samples in this study. From a to s: a, L-Aspartate; b, L-Threonine; c, L-Serine; d, L-Glutamate; e, Glycine; f, L-Alanine; g, L-Cysteine; h, L-Valine; i, L-Methionine; j, L-Isoleucine; k, L-Leucine; l, L-Tyrosine; m, L-Phenylalanine; n, L-Lysine; o, L-Histidine; p, L-Arginine; q, L-Proline; r, L-Ornithine.**

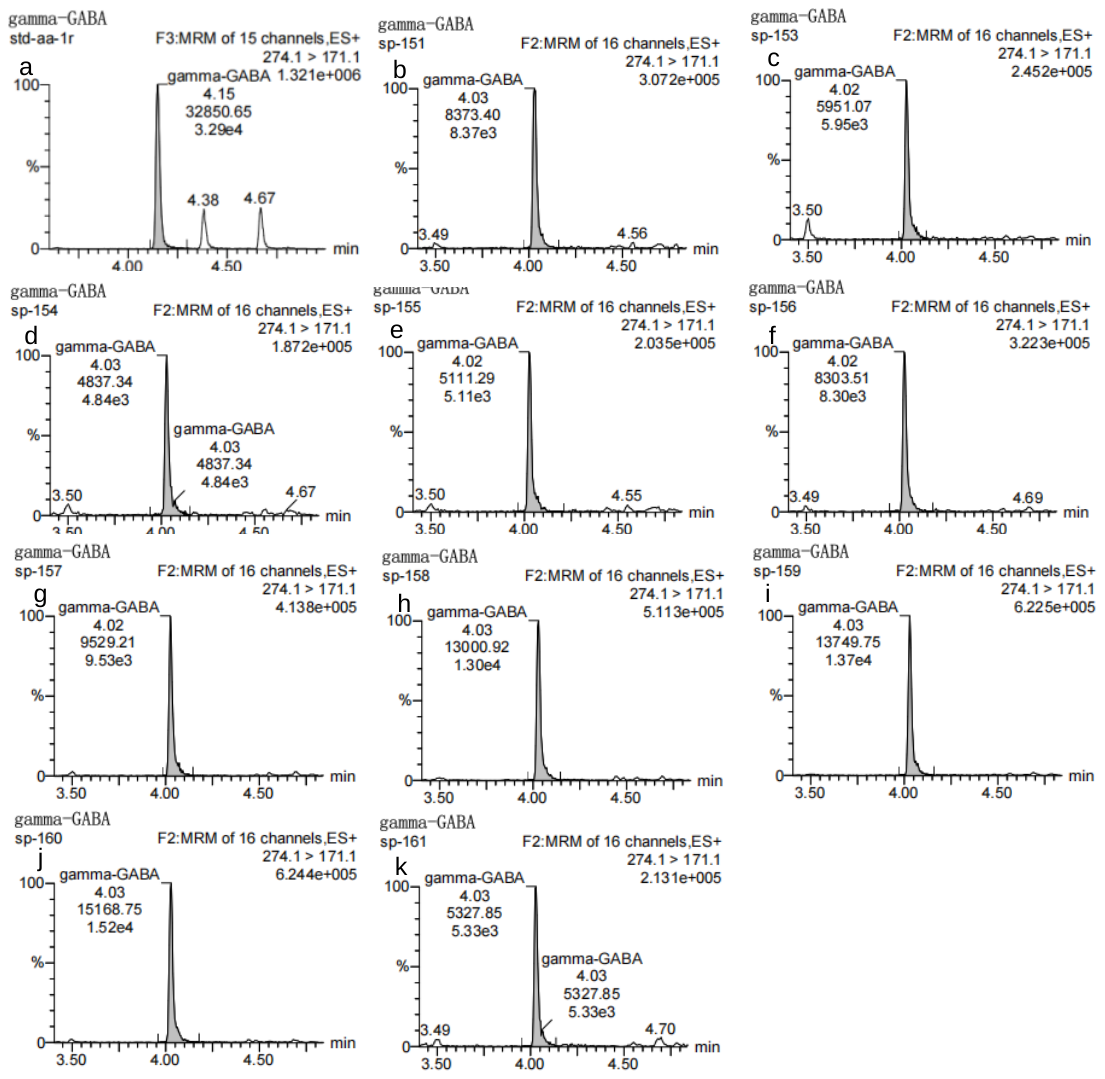

**Figure S2. Chromatogram of γ-aminobutyric acid detected in samples in this study. From a to k: a, γ-aminobutyric acid standard; b, sulfite-fueled MSA reaction network; c, sulfite-fueled MSA reaction network with addition of catalytic products from sulfite-fueled MSA reaction network; d, sulfite-fueled MSA reaction network with addition of catalytic products from sulfate-fueled MSA reaction network; e, sulfate-fueled MSA reaction network; f, sulfate-fueled MSA reaction network with addition of catalytic products from sulfite-fueled MSA reaction network; g, sulfate-fueled MSA reaction network with addition of catalytic products from sulfate-fueled MSA reaction network; h, light-Sammox-driven CRN; i, light-Sammox-driven CRN with addition of catalytic products from light-Sammox-driven CRN; j, light-Sammox-driven CRN with addition of Mg^2+^; k, light-Sammox-driven CRN with addition of Mg^2+^ + catalytic products from light-Sammox-driven CRN (Mg^2+^ addition treatment groups).**
